# Visualizing chaperone-mediated multistep assembly of the human 20S proteasome

**DOI:** 10.1101/2024.01.27.577538

**Authors:** Frank Adolf, Jiale Du, Ellen A. Goodall, Richard M. Walsh, Shaun Rawson, Susanne von Gronau, J. Wade Harper, John Hanna, Brenda A. Schulman

## Abstract

Dedicated assembly factors orchestrate stepwise production of many molecular machines, including the 28-subunit proteasome core particle (CP) that mediates protein degradation. Here, we report cryo-EM reconstructions of seven recombinant human subcomplexes that visualize all five chaperones and the three active site propeptides across a wide swath of the assembly pathway. Comparison of these chaperone-bound intermediates and a matching mature CP reveals molecular mechanisms determining the order of successive subunit additions, and how proteasome subcomplexes and assembly factors structurally adapt upon progressive subunit incorporation to stabilize intermediates, facilitate the formation of subsequent intermediates, and ultimately rearrange to coordinate proteolytic activation with gated access to active sites. The structural findings reported here explain many previous biochemical and genetic observations. This work establishes a methodologic approach for structural analysis of multiprotein complex assembly intermediates, illuminates specific functions of assembly factors, and reveals conceptual principles underlying human proteasome biogenesis.

## Introduction

A significant fraction of cellular function is carried out by large multisubunit complexes. However, due to their size and complexity, many of these complexes cannot assemble spontaneously. Instead, they are built through ordered multistep pathways orchestrated by dedicated chaperones and other assembly factors that are themselves excluded from the mature complex. One example is the proteasome, which mediates the majority of selective intracellular protein degradation and also participates in the degradation of misfolded or aggregation-prone proteins associated with neurodegenerative and other diseases.

The proteasome is a large and evolutionarily conserved eukaryotic multi-subunit complex that regulates myriad aspects of cellular function through the selective degradation of proteins. Its proteolytic active sites are located in the center of the 700-kDa barrel-shaped core particle (CP). The CP is composed of four stacked α- and β-rings (arranged α-β-β-α with 2-fold symmetry), each containing seven distinct α-subunits or seven distinct β-subunits^1,2^. The proteolytic active sites are sequestered within the CP interior and are only accessible through a narrow-gated pore formed by the α-ring at either end^3^.

Unlike conventional proteases, the proteasome harbors three different types of active site (β1/caspase-like, β2/trypsin-like, and β5/chymotrypsin-like), each of which is present twice^2,4,5^. This arrangement allows the proteasome to be both comprehensive in its substrate repertoire and processive. In addition to these features, the proteasome also shows exceptional substrate specificity. However, this specificity is not inherent in the CP, but is mostly provided through different regulators that bind the axial surface of the α-ring to open the CP gate and facilitate substrate entry. The best characterized CP regulator is the 19S regulatory particle (RP) which recognizes substrates that have designated for degradation by modification with the small protein ubiquitin^6^. Gate opening is achieved through the insertion of so-called HbYX motifs (Hb, hydrophobic; Y, Tyr; X, any residue) present at the C-termini of multiple RP subunits into the pockets that exist between adjacent α-subunits^7-14^. The monomeric CP regulator, PA200, also possesses a HbYX motif and appears to function similarly, although its cellular role remains unclear^15-17^.

Proteasomes are highly abundant. Approximately 3000 CPs are generated per minute in proliferating HeLa cells (copy number ∼4x10^6^)^18^, and their abundance can be further increased in response to stress^19^. Given this, coupled with the inherent structural complexity, it is not surprising that proteasome biogenesis depends on dedicated but transiently-associated chaperones that facilitate assembly but are not subunits of mature proteasomes (reviewed in^20-22^). Previous work has defined five CP assembly factors, proteasome maturation protein (POMP, also known as hUmp1)^23-26^ and proteasome assembly chaperones 1-4 (PAC1-4)^27-30^. In addition to these chaperones, 5 of the 7 β-subunits (β1-β2, β5-β7) are synthesized as inactive precursors with N-terminal propeptides; some of these propeptides contribute to CP biogenesis^31-34^. The importance of proper proteasome biogenesis is underscored by a group of related human diseases that arise from mutations in these assembly chaperones or regions of CP subunits that are critical for CP assembly^35-37^, and by the early embryonic lethality observed in *PAC1* knockout mice^38^.

CP biogenesis is thought to begin with the formation of a complete α-ring in a process that requires PAC1-PAC2 (hereafter PAC1/2) and PAC3-PAC4 (hereafter PAC3/4)^27-29,39-42^, with the two heterodimers bound to opposite sides of the ring. In contrast, the β-subunits are thought to incorporate sequentially onto the α-ring^33^, ultimately resulting in a half-CP, two of which then fuse to create the complete CP barrel. However, the CP remains inactive until the cleavage of the β-subunit propeptides. For the three active site subunits, this process is thought to occur through a poorly understood autocatalytic mechanism. Once activated, POMP is destroyed and PAC1/2-binding is also eliminated, resulting in mature, active 20S CP^27,43,44^.

In contrast to PAC1/2, PAC3/4 are released early during CP biogenesis, although the timing and trigger for release have remained unclear. PAC3/4 are not present in the best-characterized assembly intermediate, the 13S complex, which has been identified even in wild-type mammalian cells and which consists of PAC1/2, the complete α-ring, and β2-4^34,45-47^. Although no structural information is yet available for PAC3/4-containing intermediates, modelling of an isolated yeast α5-Pba3-Pba4 crystal structure onto mature CP suggested steric clash between Pba3/4 and the β4 subunit, leading to the notion that insertion of β4 might induce release of Pba3/4^39^. Alternately, knockdown of β3 in human cells resulted in accumulation of an intermediate containing all 5 chaperones^33^, raising the possibility that PAC3/4 release is coordinated with insertion of β3, despite the lack of obvious steric clash. Still a third possibility was suggested by high-resolution structures of yeast 13S and pre-15S intermediates which revealed major steric clash between Pba3/4 and Ump1^48^.

In addition to this uncertainty regarding PAC3/4, other major aspects of CP biogenesis remain poorly understood. Most notably, these include the mechanisms that account for the highly regimented stepwise production of the β-ring and the mechanism that couples CP assembly with peptidase activation^31,43,44,49^. Here, we have used cryo-EM to obtain and characterize the first structures of human assembly chaperone-containing proteasome core particle subcomplexes. We report a structure of the first PAC3/4-containing CP intermediate, as well as a series of structures containing increasing numbers of β-subunits, yielding detailed understanding of the stepwise assembly of the β-ring. This study significantly advances our understanding of proteasome assembly, with implications for the broader field of chaperone-mediated multiprotein complex biogenesis.

## Results

### A system for recombinant human CP expression

Recombinant systems have proven useful for studying mature proteasome complexes ^16,50-52^. To express the human CP in insect cells, we utilized a combination of the biGBac^53^ and MultiBac^54^ systems to generate baculovirus transfer vectors: one with all seven α-subunits, another with all seven β-subunits, and the third with the five CP assembly chaperones. To allow for affinity purification, we appended a C-terminal Twin-Strep tag to either β2 or β7, the first and last CP subunits, respectively, to incorporate. We affinity purified CP, followed by size exclusion chromatography (SEC). Analysis of both preparations (Fig. 1a-b) confirmed the expected electrophoretic properties for mature CP. Furthermore, cryo-EM analysis of β2-tagged purified CPs yielded a high-resolution structure that superimpose with native human 20S CP (RMSD 0.703Å) (Fig. 1c right panel, Extended Data Fig. 1).

**Fig. 1.**
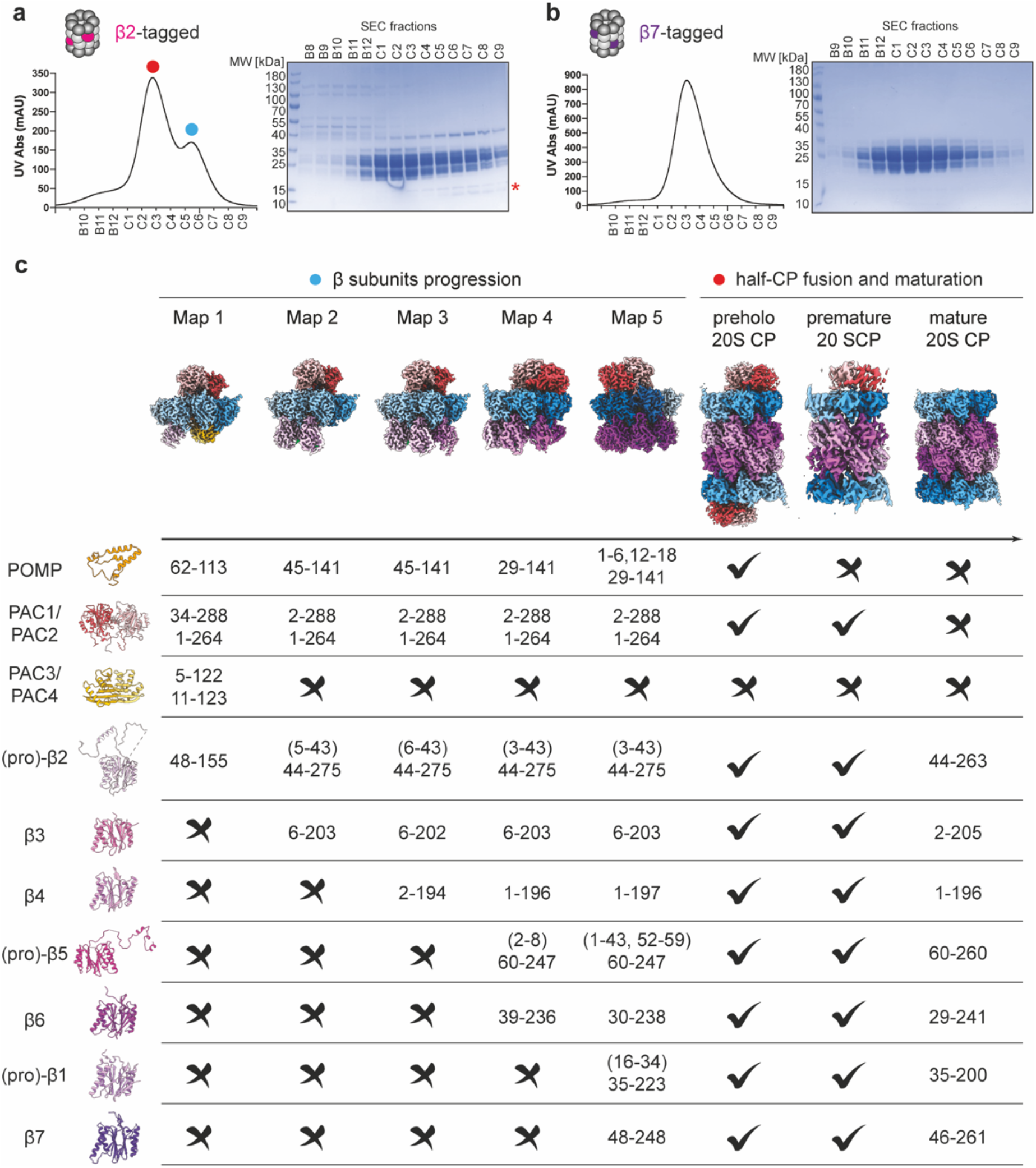
Recombinant expression of human 20S proteasome in insect cells. **a**, Purification of recombinant β2-tagged (PSMB7) 20S CP. Eluates were subjected to size exclusion chromatography (SEC) (right panel), and the fractions analyzed by SDS polyacrylamide gel electrophoresis (SDS-PAGE) followed by Coomassie staining (left panel). Asterisk indicates novel bands in the later fractions that likely correspond to the assembly chaperones, PAC1-4 and POMP. Red dot indicates main peak fractions corresponding to half CP fused 20SCP and blue dot indicates shoulder peak fractions corresponding to 20SCPassembly intermediates. **b**, Purification of recombinant β7-tagged (PSMB4) 20S CP. Eluates were subjected to size exclusion chromatography (right panel), and the fractions analyzed by SDS polyacrylamide gel electrophoresis (SDS-PAGE) followed by Coomassie staining (left panel). **c**, Cryo-EM maps of eight distinct CP subcomplexes. Their subunits, chaperone configurations, and the resolved region in each map are indicated.

The SEC profile for the β2-tagged CP revealed a shoulder peak consistent with lower molecular weight subcomplexes (Fig. 1a). Analysis of these fractions by SDS-PAGE revealed additional proteins 12-15 kDa in size. Some assembly chaperones would be expected to migrate in this size range, suggesting that these complexes might represent immature CP species. We pooled and analyzed these fractions by cryo-EM, yielding structures of five distinct immature CP subcomplexes with overall resolutions ranging from 2.67 to 2.95 Å (Fig. 1c left panel, Extended Data Fig. 2, Supplementary Table 1). We order these structures based on the number of visible β-subunits, with their compositions indicated in Figure 1c. We present the structures with the α-subunits in blue and the β-subunits in purple, increasing in shade by subunit number (α-ring) or the order of incorporation (β-ring). These structures are fully consistent with the ordered, sequential addition of β subunits proposed from genetic knock-down studies^33^ in human cells. Structure 1 represents the earliest CP subcomplex visualized to date from any organism and consists of the α-ring, all five chaperones, and β2 (Figure 1c, Supplementary video 1). Structures 2-5 consist of the α-ring, chaperones POMP and PAC1/2, and an increasing number of β-subunits (Figure 1c). Structures 3 and 4 are analogous to recent structures of 13S and pre-15S proteasome assembly intermediates, respectively, determined from yeast^48^. Thus, the structural progression of human complexes lies within a conserved proteasome assembly pathway. In addition to mature CP, detailed analysis of the cryo-EM data from the higher molecular weight fraction revealed two supra-20S structures referred to as preholo-20S CP and premature-20S CP (Fig. 1c right panel, Extended Data Fig. 1d-f). Comparing these seven different structures to each other and to the matched mature CP allows for visualization of conformational progression across the assembly pathway and the specific molecular interactions occurring at each step.

### Structure 1: α-ring stabilization and β-ring initiation through coordinated chaperone function

Structure 1 shows PAC1/2 and PAC3/4 within the same complex for the first time (Fig. 2a-b, Supplementary video 1), and illustrates POMP-dependent initiation of β-ring assembly through its binding to both the α-ring and β2 (Figure 2d). Together PAC1/2 and PAC3/4 bind all 7 α-subunits with PAC1/2 perched atop the α-ring on the same surface recognized by proteasome activators. This placement appears to serve two main purposes. First, it likely helps coordinate α-ring configuration since PAC1/2 contacts all α-subunits except α3 (Fig. 2b bottom panel). Second, as observed for its yeast counterpart Pba1/2 in later proteasome assembly intermediates^21^, PAC1/2 directly binds multiple α-subunit N-termini and, through numerous interactions, orchestrates an open-gate conformation (Fig. 2b). Perhaps most dramatically, α5’s N-terminus was extensively bound by PAC1/2 and was rotated upward, extending more than 20 Å from the α-ring (Fig. 2a, Supplementary video 1). This interaction likely contributes to PAC1/2’s previously reported ability to bind α5 in isolation^27^.

**Fig. 2.**
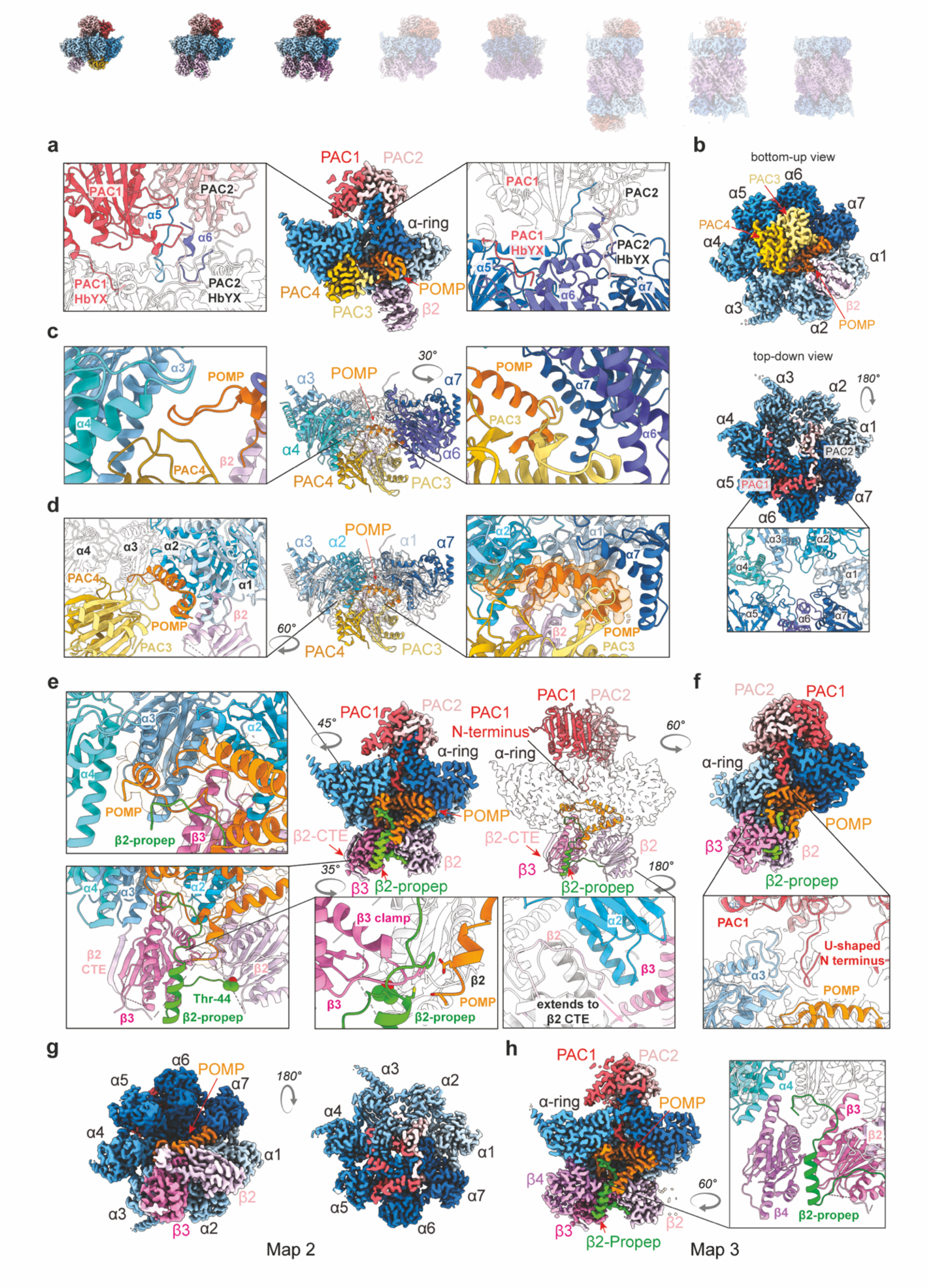
Structural progression of early assembly intermediates. **a**, Cross-section view of cryo-EM map of the structure 1 shows all five assembly chaperons. PAC1/2 is perched on top of the α-ring and interact with all α-subunits except for α3. **b**, Bottom-up view (top panel) of cryo-EM map of structure 1 shows that PAC3/4 resides in the groove between α4/5 and α5/6, and β2 sits in the groove between α1 and α2. Top-down (bottom panel) of structure 1 shows that PAC1/2 holds the α-ring ‘gate’ open and the pore is open. **c**, Side view of structure 1 shows that PAC3/4 interacts with α3-α7 of α-ring. **d**, Side view of structure 1 shows that POMP interacts with α1-α3 and α7 of α-ring. **e**, Cross-section view of cryo-EM map of structure 2 shows incorporation of the β3 is mutually stabilized by β2-propeptide and β2 C-terminal extension (CTE). Close-up view shows additional N-terminal region of POMP interacting with β2-propeptide and β3. **f**, Close -up view of PAC1 N-terminus of structure 2 shows its interaction with α3 and POMP. **g**, Bottom-up view (left panel) of cryo-EM map of structure 2 shows that release of PAC3/4 frees up α4/5 and α5/6 grooves and β3 occupies α2/3 groove. Top-down view (right panel) shows that the pore is closed. **h**, Cross-section view of cryo-EM map of structure 3 shows incorporation of the β4 and its interaction with β3 and the α-ring.

The visualized N-termini of the α-subunits largely resemble those seen in substrate-engaged 26S proteasome^55^ (Extended Data Fig. 3a). Thus, the arrangement in Structure 1 appears poised to allow threading of polypeptides through the α-ring pore. This strikingly contrasts from mature isolated CP, where α-subunit N-termini extend into and block access to central pore^2^. Some proteasome activators trigger gate opening via their C-terminal HbYX motifs that insert into pockets between adjacent α-subunits^10,11,15-17,56^. PAC1 contains a conserved, canonical HbYX motif (Ile-Tyr-Thr), which is inserted into the α5/α6 pocket (Fig. 2a). PAC2 contains a variant sequence (Leu-Phe) and is inserted more shallowly into the α6/α7 pocket, with Phe positioned where Tyr sits in more conventional HbYX motifs (Fig. 2a).

PAC3/4 sits on the opposite face of the α-ring (i.e., the site of the future β-ring). Consistent with a prior crystal structure of the orthologous isolated yeast Pba3/4-α5 complex^39,57,58^, PAC3/4 occupy the α3-α7 side of the ring and, interestingly, a portion of α5 is situated at the PAC3/4 interface (Fig. 2b-c, Extended Data Fig. 3b). PAC3/4’s position on the α-ring corresponds to that of the β4-β7 subunits later during assembly (Extended Data Fig. 3c). By occupying these sites, PAC3/4 not only stabilize the α-ring to provide a platform for β-ring assembly, but also preclude incorporation of the late β-subunits, helping to set the order of β-subunit incorporation.

Unexpectedly, PAC3/4 also makes multiple contacts to the β-ring chaperone POMP. Interestingly, the same site in POMP bound by PAC4 is recognized later during assembly by the β5 propeptide (Extended Data Fig. 3f). As described below, POMP consists of a series of helices and loops that become progressively more resolved upon successive β-subunit addition, with POMP winding back and forth between the α- and β-rings. Here, one end was inserted into the groove between PAC3, α7, and α1, with the other end inserted between PAC4, α3 and α4 (Fig. 2d). These contacts rationalize previous observations regarding POMP’s direct interaction with the α-ring in isolation and its interaction with α3, α4, and α7 by yeast two hybrid analysis^33,59^.

Finally, Structure 1 reveals the basis for β-ring initiation with β2: a PAC3/4-bound complete α-ring engages POMP, which in turn can recruit β2. A portion of β2 is visible in Structure 1, inserted between α1 and α2, and in contact with POMP (Fig. 2d). At this stage of assembly, β2 is poorly resolved, presumably due to the lack of stabilization by adjacent β-subunits.

### Role of the β2-propetide and POMP in the sequential addition of β3 and β4

Structure 2 is distinguished by the loss of PAC3/4 and the addition of β3 (Fig. 2e, Supplementary video 2). Several features of the structure explain previous genetic observations from mammalian cells that β3 is the second β-subunit and requires prior β2 addition. The β2 subunit grasps β3 with both a C-terminal extension (CTE) and the β2 propeptide (Fig. 2e), explaining the requirement for both regions to properly incorporate β3 in mammalian cells^33^. Extending from the β2 active site threonine, the propeptide meanders, forms a helix crossing the interior surface of β3, and extends to terminate in a loop that secures newly visible regions of POMP that co-fasten β3 in the groove between α2 and α3 (Fig. 2e, bottom-left panel). Meanwhile, part of the β2 CTE forms a strand that extends a β-sheet from the adjacent β3, an arrangement also seen in mature CP^2,60^.

β3 is stabilized by direct interactions with POMP, as reflected in newly visible residues of POMP (residues 45-62 and 114-141) in Structure 2 compared to Structure 1(Fig. 2e). Residues 114-141 directly clamp β3 against the α-ring through an interaction that is also buttressed by part of the β2-propeptide (Fig. 2e, bottom panel). Meanwhile, POMP residues 45-61 buttress the adjacent portion of the β2-propeptide that secures the interior surface of β3. Notably, the regions of the β2-propeptide and POMP that clamp β3 in place bind to the same α3/α4 surfaces as PAC3/4 did in Structure 1 (Fig. 2e, left panel), suggesting a potential mechanism for the eviction of PAC3/4 based on the stabilization of those regions of POMP and the β2 propeptide upon incorporation of β3. Together this could explain why knockdown of β3 causes accumulation of the five chaperone-containing intermediate chaperones^33^ without β3 directly overlapping with PAC3/4.

With the departure of PAC3/4, the stability of the α-ring may be further supported not only through additional POMP interactions that bridge β2/β3 and the α-ring, but through changes at the α-ring pore. The N-termini of α1 and PAC1 are newly observed in Structure 2, passing through the gate towards POMP (Fig. 2e-f, Supplementary video 2). PAC1’s N-terminus runs alongside α1’s N-terminus through the center of the α-ring until PAC1 contacts a newly ordered region of POMP and makes a U-turn to terminate in the center of the α-ring pore (Fig. 2f-g). Concomitantly, additional residues of the gating N-termini of α2-α4 are visible, stabilized in an open conformation by a collar of Tyr residues between α-ring subunits rather than by direct, extensive interactions with PAC1/2 as is seen for α5 (Extended Data Fig. 6). This open gate but occluded pore arrangement is maintained throughout the other assemblies we describe including the preholo-20S CP.

Key differences between Structures 1 and 2 set the stage for incorporation of β4 seen in Structure 3 (Fig. 2h). One side of β4 binds a composite surface composed of β3 and the β2 propeptide, while its other side binds the region of the α4 that was previously bound to PAC4 in Structure 1 (Fig. 2h, right panel, Extended Data Fig. 3c, Supplementary video 3). Importantly, Structure 3 corresponds to the well-recognized 13S precursor^34,45^, and is largely superimposable on the yeast 13S^48^, although there are some notable species-specific differences. First, while the N-termini of both PAC1 and yeast Pba1 are threaded through the open gate into the CP interior, in yeast it is the N-terminus of α2^56^, (not α1) that runs alongside Pba1’s N-terminus into the CP interior (Extended Data Fig. 3e). Second, PAC1’s longer N-terminus loops back upwards towards the α-ring gate whereas Pba1’s ends in contact with Ump1 and the β5-propeptide^48^ (Extended Data Fig. 3e).

### Establishment of a POMP/propetide network accompanies completion of the β-ring

Structure 4 corresponds to the previously reported yeast pre-15S intermediate^48^ and contains the α-ring, β2-6, PAC1/2, and POMP (Supplementary video 4), while Structure 5 contains β1 and β7 and shows better resolution of β6. Together, these structures illustrate how portions of assembly factors become progressively structured as they secure remaining β-subunits in a stepwise manner (Extended Data Fig. 5). Insertion of β5 and β6 is accompanied by increased resolution at the N-terminus of POMP (residues 29-44) (Fig. 3a,c), which is accompanied by redirection of the N-terminus of the β2 propeptide to fill a newly formed pocket at the confluence of POMP, β4-5, and α3-4 (Fig. 3b,d).

**Fig. 3.**
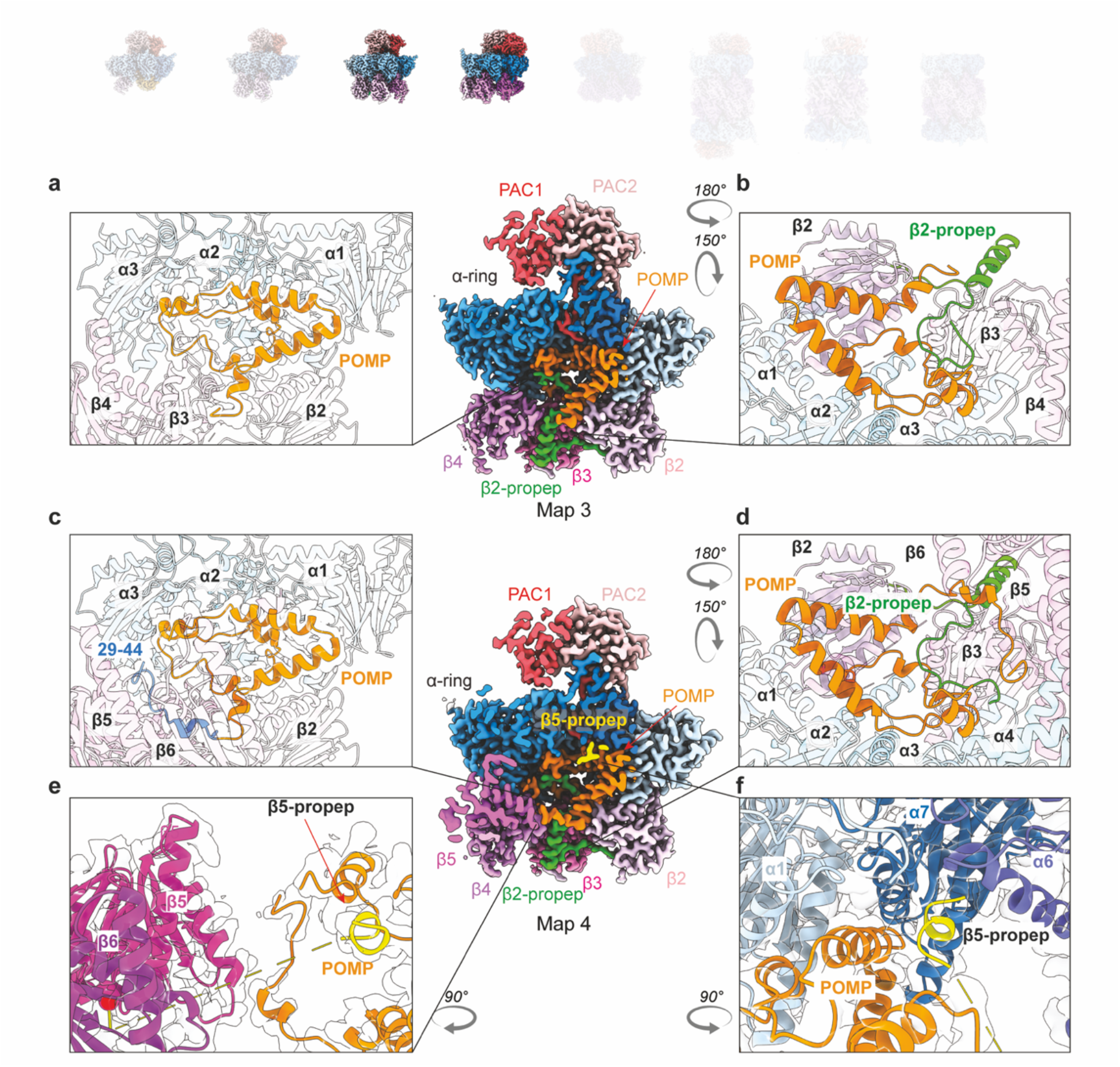
Distinct structural features upon β5 and β6 incorporation. **a-b**, Cross-section view of cryo-EM map of the structure 3 shows PAC1/2, POMP and β2-β4. Close-up view of POMP (**a**) shows its helix-turn-helix configuration and its interaction with the α-ring and β-subunits. Close-up view (**b**) reveals interaction between POMP and β2-propeptide. **c-f**, Cross-section view of cryo-EM map of the structure 4 shows PAC1/2, POMP and β2-β5 (β6 is cropped in the cross-section) upon β5 and β6 incorporation. Close-up view of POMP (**c**) shows that additional N-terminal region of POMP (blue) is resolved and stabilizes β5 and β6 incorporation. Close-up view of β2-propeptide (**d**) reveals the additional resolved N-terminal region of POMP interacts with the N-terminus of β2-propeptide and drives it to a different orientation. Close-up view of β5, β6 and POMP (**e**) shows that the additional resolved N-terminal region of POMP interacts with β5 loop. Close-up view of β5-propeptides (**f**) shows its interaction with α6, α7, α1 and POMP.

In Structure 4, only the extreme N-terminus (residues 2-8) of the β5 propeptide is visible at the junction of α6, α7, and POMP, resembling a harpoon as it anchors 60Å away from the rest of β5 (Fig. 3e-f). Structure 5 shows the ultimate arrangement of assembly factors within a completed β-ring just prior to half-CP fusion (Supplementary video 5). Here, additional elements in the β5 propeptide and POMP visibly form a belt supporting the β5-β1 subunits (Fig. 4a-d). Regions of the β5 propeptide between the N-terminal anchor and the catalytic domain are newly visible in Structure 5 compared to Structure 4, suggesting that its folding is coordinated with progressive addition of β-subunits (Fig. 4a,c, Extended Data Fig. 5). Tracing the β5 propeptide from where it emerges from β5, it winds back and forth between the β-ring and α-ring to secure β6, β7, and β1, and crosses POMP before terminating with the interactions already observed in Structure 4 (Fig. 4d,f). The extensive interactions of the β5 propeptide at β6’s interface with α-subunits clarify observations that the β5 propeptide is essential for the incorporation of β6, while β5 does not require its own propeptide for incorporation^33^. Meanwhile, newly visible N-terminal regions of POMP in Structure 5 support the β6-β5 and β1-β2 pairs of subunits. A portion of the β1 propeptide (residues 16-34) is also visualized for the first time in Structure 5 and resembles a staple extending from β1 to β7 (Fig. 4e).

**Fig. 4.**
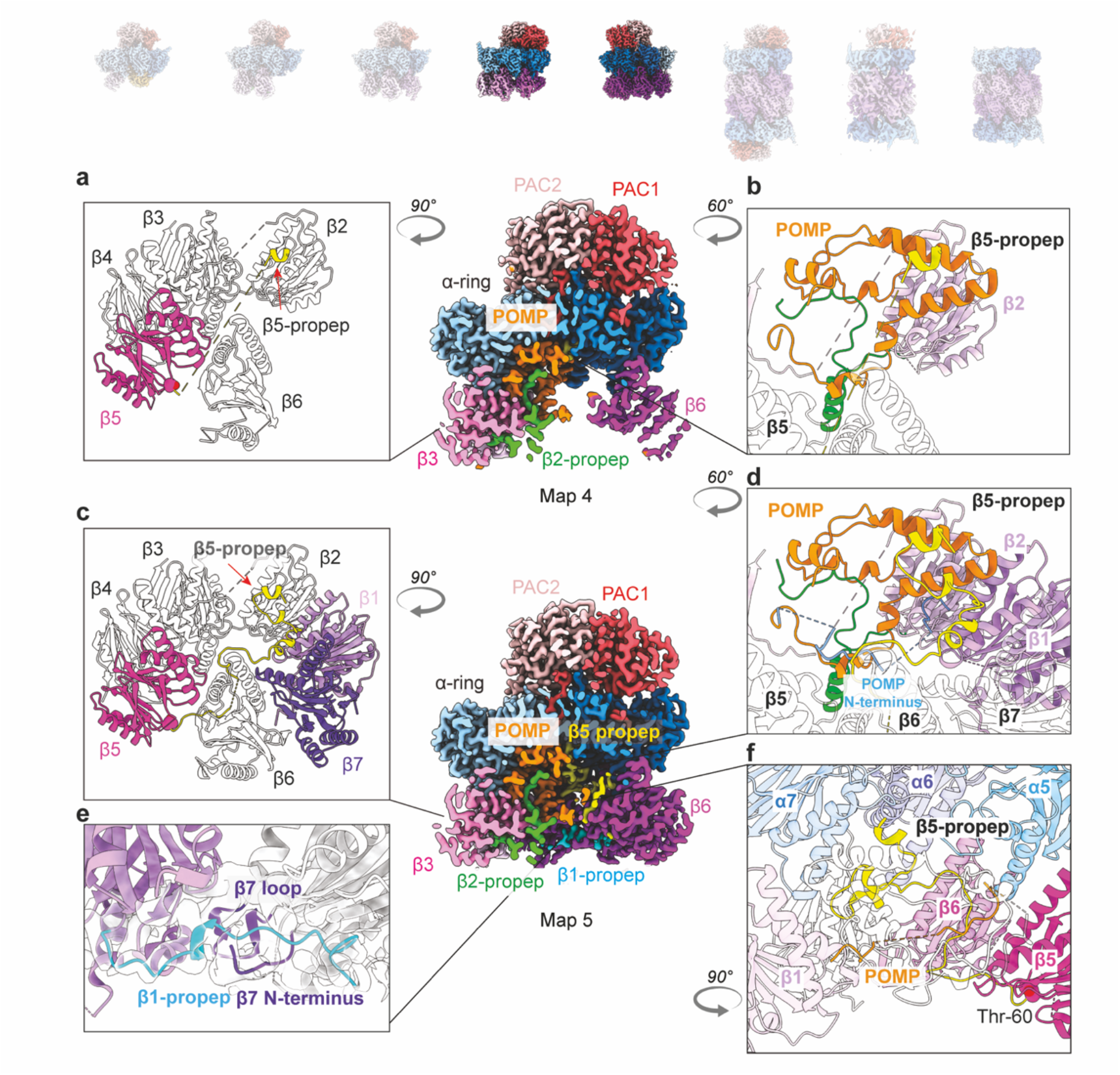
Incorporation of β7 and β1 completes β-ring and reveals all β-propeptides. **a-b**, Cross-section view of cryo-EM map of the structure 4 shows PAC1/2, POMP, β3 and β6 (β2 is hidden behind and β4-5 are cropped in the cross-section) upon β5 and β6 incorporation. Close-up view of β-ring (**a**) and interaction between β5-propeptide and POMP (**b**). **c-f**, Cross-section view of cryo-EM map of the structure 5 shows PAC1/2, POMP, β3 and β6 (β7 and β1-2 are hidden behind and β4-5 are cropped in the cross-section) upon β7 and β1 incorporation. Close-up (**c**) shows additional β5-propeptide is resolved upon completion of the β-ring (blue) compare to β5 in Structure 4. Close-up view of β5-propeptide’s interaction with POMP, β1 and β2 (**d**) shows that POMP’s N-terminus (blue) is resolved upon completion of the β-ring and interacts with β5-propeptide. Close-up view of newly incorporated β1 and β7 (**e**) reveals the β1-propeptide and its interaction with β7 N-terminus and loop. Close-up view (**f**) of β5-propeptide’s interaction with α5-7, β5-6 and β1.

### Half-CP fusion and proteolytic activation

Comparing the half-CP from Structure 5 with the preholo-, premature- and mature 20S CP structures reveals conformational changes accompanying half-CP fusion and subsequent loss of propeptides, POMP, and PAC1/2 to form the mature 20S proteasome (Fig. 5a). With the notable exception of the β6 subunit, which becomes largely visible in Structure 5 and superimposes with the post-fusion structures, the most striking differences between Structure 5 and preholo-20S CP are in conformations of the β-subunits, and show how fusion of two half-CPs coaxes the proteases towards active conformations (Fig. 5b-c). In particular, two key elements from each subunit along the β-ring/β-ring interface of post-fusion CP structures are a β-hairpin (e.g., residues 61-75 of β2) and adjacent loop (e.g., residues 205-215 of β2) (Fig. 5d). We refer to these as “fusion-hairpin” and “fusion-loop”, respectively, because these interlock in a zipper-like structure with those from the opposing subunit across the β-ring/β-ring interface. However, Structure 5 shows that in the half-CP, these elements are exposed and the majority are either not visible and presumably dynamic (the β2, β3, β5, and β1 subunits), or the β-hairpin is splayed relatively outward or inward toward the center of the subunit (the β4 and β7 subunits) (Extended Data Fig. 4). The fusion-hairpin and fusion-loop become visible and/or are reoriented upon fusion as seen in the preholo intermediate, where they adopt conformations matching those in the mature CP (Fig. 5d-g, Extended Data Fig. 4).

**Fig. 5.**
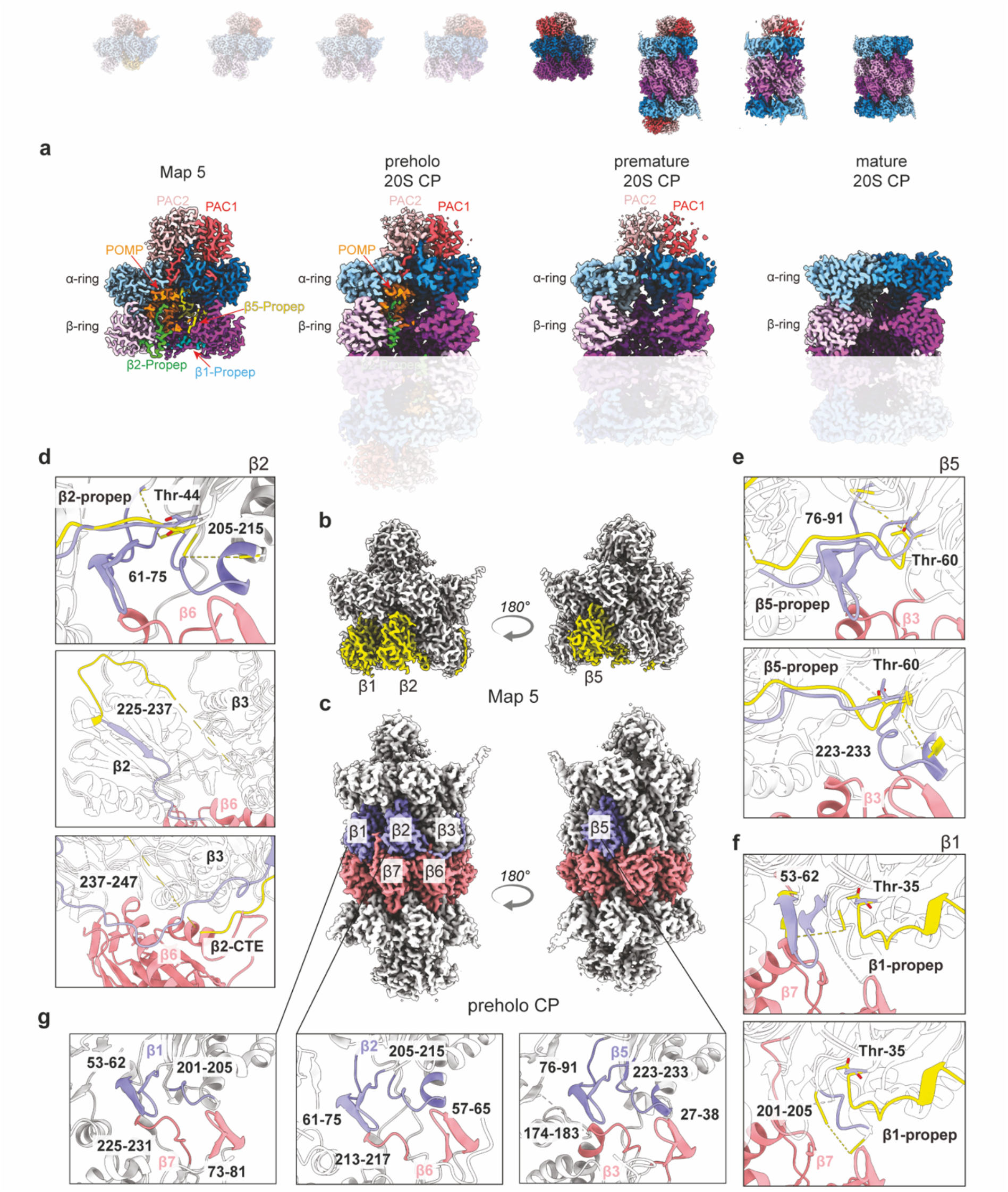
Half CP fusion and 20SCP maturation. **a**, Cross-section view of cryo-EM map of structure 5, preholo-20SCP, premature-20SCP and mature 20SCP shows the presence and release of assembly chaperones and β-propeptides during 20SCP fusion and maturation process. **b**, Cryo-EM map of structure 5 with β1, β2 and β5 colored in yellow. **c**, Cryo-EM map of preholo CP with β1, β2 and β5 of one half CP colored in purple and the other half CP colored in salmon. **d**, Close-up view of β2 structural changes (β2 from structure 5 colored in yellow and β2 from preholo CP colored in purple) and its interaction with β6 (salmon) from the opposite half CP upon CP fusion. **e**, Close-up view of β5 structural changes (β5 from structure 5 colored in yellow and β5 from preholo CP colored in purple) and its interaction with β3 (salmon) from the opposite half CP upon CP fusion. **f**, Close-up view of β1 structural changes (β1 from structure 5 colored in yellow and β1 from preholo CP colored in purple) and its interaction with β7 (salmon) from the opposite half CP upon CP fusion. **g**, Close-up view of fusion-tetrad formed by the fusion-hairpin and fusion-loop from the opposing β subunits.

Overall structural similarity in the β1, β2, and β5 subunits in preholo intermediate and mature 20S CP is consistent with the notion that fusion plays a major role in driving catalytic activation of the proteasome. When ordered post-fusion, the fusion-hairpin and fusion-loop abut the catalytic Thr residue of β1, β2, and β5, and shape the active sites of the protease subunits. Furthermore, the fusion-loop contains key residues required for autocatalytic protease activation^44^.

For the β2 subunit, there is additional, major restructuring in the protease domain. In Structures 3-5, i.e., from the point in β-ring assembly when β2 was largely visible, β2’s residues 225-237 face the α-ring and cling to adjacent α2 subunit. This interaction was not observed in proteasome assembly intermediates from yeast due to its substantially different sequence and structure of the α2 subunit in this vicinity (Extended Data Fig. 3d). For human β2, the post-fusion structures show residues 225-237 making a U-turn and proceeding 140° in the opposite direction, towards the other half CP (Fig. 5d, middle panel). This structural remodeling completes the β2 subunit’s protease fold, commensurate with the fusion of two half CPs. This conformational change also directs the ensuing β2 residues (237-247) towards the opposite half-CP (Fig. 5d, bottom panel). These β2 residues comprise the portion of the CTE that connects the β2 protease domain and C-terminal strand bound to β3 and that was not visible and presumably dynamic the structures across β-ring assembly. In the post-fusion structures, this intervening region of the β2 CTE forms a belt that engages β6 from the opposite half-CP.

Interestingly, in the preholo CP structure, density corresponding to the propeptide is continuous with the β2 and β5 active sites (Fig. 5d,e, Extended Data Fig. 4h-j). Meanwhile, prior crystallographic and functional analyses of yeast proteasomes harboring propeptide and active-site adjacent mutations suggested that subtle perturbation of (1) the backbone conformation at the junction between the active site and propeptide, (2) placement of adjacent side-chains including a critical lysine, and (3) the constellation of surrounding water molecules can influence the rate of propeptide cleavage^31,44^. Although the resolution of our structures precludes precise placement of atoms and visualization of waters, the preholo CP intermediate structure shows features poised to impact these facets of propeptide maturation. First, the β2 and β5 propeptides make numerous contacts that impact their orientation. For the β2 propeptide, these mirror those in Structure 5, and include interactions with β3, β4, α3, and POMP (Fig. 5d). Meanwhile, 52-59 residues of the β5 propeptide are observed retaining contacts with β6 (Fig. 5e). Second, the propeptides themselves align the active site and contact adjacent residues including the lysine that is critical for their cleavage^44^ (Fig. 5d,e, Extended Data Fig. 4h-j).

In the preholo-20S structure, both PAC1/2 and POMP remain bound to the α and β-rings (Fig. 5a). POMP in both half CPs is observed in the middle of the CP interior, and it maintains its interaction with β1-5 including the β2-propeptide. POMP is not visible in the premature-20S, suggesting that cleavage of the propeptides might also destabilize these interactions, allowing for its degradation. Persistence of at least one PAC1/2 dimer in the premature-20S structure suggests that PAC1/2 elimination from the complex occurs after POMP degradation, and therefore they are poised to maintain the open gate conformation until the very end of CP maturation.

## Discussion

Given their central role in cellular regulation, it is important to understand how proteasomes are generated (Fig. 6). The seven cryo-EM reconstructions of recombinant human CP assembly complexes reported here include the first structures from any species for several proteasome assembly intermediates and the first structures containing PAC3/4. Structure 1 reveals how the β-ring is initiated and why the first subunit is confined to β2. This structure also reveals two probable and unexpected contributions of a so-called α-ring chaperone to the β-ring. First, PAC3/4 may help to recruit POMP through direct interaction and, second, PAC3/4 may help order β-subunit incorporation by preventing premature incorporation of the later subunits through occlusion of their binding sites on the α-ring. Structure 2, when compared with Structures 1 and 3, shows for the first time progressive steps by which individual subunits are added. Structure 5, the half-CP, reveals the function of the β5 propeptide in completing β-ring assembly and its collaboration with POMP in this process; Structure 5 also shows how the proteasome active sites are restrained prior to fusion with another half-CP.

**Fig. 6.**
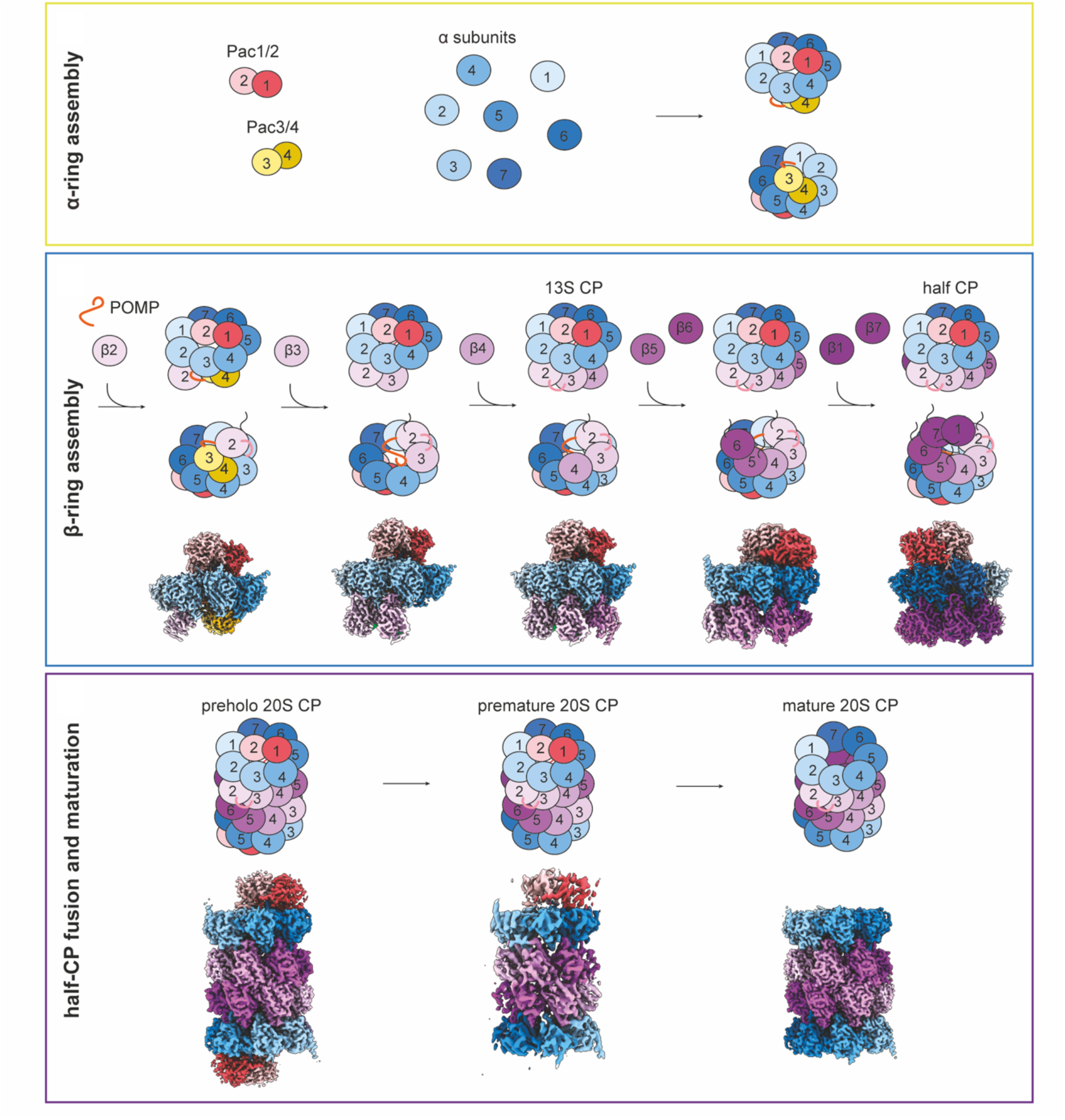
Schematic representation of chaperon-mediated stepwise assembly of the 20S CP and summary of high-resolution structures of human assembly intermediates. Assembly of the 20S core particle can be distinguished into the three steps: (i) α-ring assembly, (ii) β-ring assembly, and (iii) half CP fusion and maturation. High resolution maps of human CP assembly intermediated presented in this study are indicated below their schematic representations

Notably, Structures 3, 4, and the human preholo intermediate also allow comparison to the previously-determined structures of the corresponding yeast 13S, pre-15S, and preholo intermediates^48,61^, not only validating our approach but also revealing conserved and species-specific aspects of proteasome assembly. One particularly striking feature was the overall high degree of structural conservation of POMP, the β2-propeptide, and the β5-propeptide, respectively, despite relatively low sequence similarity between the two species. The structures also reveal key conceptual principles concerning the generation of directionality during CP biogenesis, which may have relevance for other molecular machines. In principle, the assembly of any 28-subunit complex could proceed through an almost infinite number of distinct pathways. In the mature CP, each subunit is engaged in lateral, vertical, and even diagonal interactions with its neighbors. These interactions are reflected in the exceptional stability of the CP, which can withstand harsh purification conditions (e.g., 0.5 M salt washes)^48,62^. These interfaces cannot be fully occupied during CP assembly. Many unpartnered regions of proteasome subunits and assembly factors appear flexible, as reflected in the lack of density in the cryo-EM maps. Each incoming subunit “structures” or rigidifies its neighboring subunit or chaperone in a templating-like manner. Because each interface is unique, binding affinity should naturally be highest for true neighboring subunits, and this should drive the pathway forward through energetic favorability. Insertion of each new subunit not only stabilizes the existing assembly intermediate, but also promotes the formation of the next intermediate until the β-ring is complete. It would seem that greatly reducing the number of available assembly pathways would ensure a high degree of uniformity and fidelity in CP biogenesis.

Interestingly, it seems that premature incorporation of subunits is also prevented by a series of previously unknown structural checkpoints. For example, the chaperones initiating β-ring assembly -PAC3/4 and POMP - together bind all α-subunits. This ensures that β-ring assembly is initiated only after completion of the α-ring. As another example, several structural features require proper placement of β2-β4 prior to β5 incorporation. Thus, the β6, β1, and β7 subunits, which are progressively secured by the β5 propeptide, are stabilized only after insertion of β5.

Another important aspect of CP assembly relates to the principal role of the active site subunit propeptides in helping recruit and stabilize their neighboring subunits. The propeptides appear not to act in isolation, but through a complex network of interactions with POMP. Indeed, one can conceive of CP assembly as a process of progressively building an internal structured scaffold within the growing complex, and then self-disassembling that scaffold once activation has occurred. Interestingly, many higher organisms encode additional proteasome β-subunits which are incorporated into immune-specific CP complexes^20^. These subunits show considerable sequence divergence in their propeptides, and it will be interesting to determine the molecular consequences of this arrangement.

Similar to β-ring assembly, but on a much larger scale, complementary protein segments not observed in the chaperone-bound half-CP (Structure 5) simultaneously become visible in the preholo intermediate. These include what we refer to as the fusion-hairpins and fusion-loops, which in β1, β2, and β5 configure the active sites including during autocatalytic propeptide processing^44^. Also, it is intriguing that the one subunit that shows little structural change between the half-CP and preholo intermediate is β6. Its midline partner, β2, is the one that apparently undergoes the most large-scale structural remodeling during fusion of two half-CPs. It is tempting to speculate that β6 may assist in templating this process, with its pre-formed fusion-hairpin and fusion-loop capturing those from an opposing β2 subunit. It is possible that β6’s ensnaring the opposite β2 CTE loop acts as a lever that redirects the preceeding sequence and promotes completion of the adjacent protease fold.

This work has important clinical implications as defects in CP assembly are responsible for a rapidly growing family of related diseases characterized by immune dysregulation^21,35-37^. Our study provides a system for future studies aimed at determining functional consequences of individual mutations, including patient-derived mutations, on CP assembly. In the long-term, a more complete understanding of CP assembly in the normal and diseased states might even lead to therapeutic interventions to enhance or impair the efficiency of CP biogenesis, as needed clinically. Finally, we believe that strategy employed here, namely to visualize sequential intermediates along complex assembly pathways, could be applied to many other molecular machines, which in turn could be helpful in understanding diseases characterized by defects in multiprotein complex biogenesis.

## Materials and Methods

### Cloning, Protein Expression, and Purification

The cDNAs encoding 20S CP subunits and assembly chaperones were either obtained from an in-house human cDNA library (Max Planck Institute of Biochemistry) or ordered as synthetic cDNA from Eurofins or IDT, if multiple restriction sites were to be occluded. Baculovirus transfer vectors were assembled by utilizing a combination of the biGBac system^53^ and 3^rd^ generation MultiBac system^54^. In short, all subunits were first cloned by Gibson assembly into the vector pACEBac1 which was utilized as a library vector. For β2 (PSMB7) and β7 (PSMB4), both untagged and TEV-cleavable C-terminal twin strep tag constructs were constructed. Primers for step 1 assembly of the biGBac system were modified to prime before the polh promoter and SV40pA site within pACEBac1, respectively. Step one assembly reactions were set up to obtain the vectors pACEBac1-POMP-PAC1-PAC2-PAC3-PAC4 (Addgene plasmid #XXXX), pACEBac1-PSMA1-PSMA2-PSMA3-PSMA4, pACEBac1-PSMA5-PSMA6-PSMA7, pACEBac1-PSMB1-PSMB2-PSMB3-PSMB4, pACEBac1-PSMB1-PSMB2-PSMB3-PSMB4-TEV-2xSTII, pACEBac1-PSMB5-PSMB6-PSMB7, and pACEBac1-PSMB5-PSMB6-PSMB7-TEV-2xSTII. Subsequently, the final vectors pACEBac1-PSMA1-PSMA2-PSMA3-PSMA4-PSMA5-PSMA6-PSMA7 (Addgene plasmid #XXXX), pACEBac1-PSMB1-PSMB2-PSMB3-PSMB4-TEV-2xSTII-PSMB5-PSMB6-PSMB7 (Addgene plasmid #XXXX), and pACEBac1-PSMB1-PSMB2-PSMB3-PSMB4-PSMB5-PSMB6-PSMB7-TEV-2xSTII (Addgene plasmid #XXXX) were assembled via restriction cloning utilizing the MultiBac multiplication module.

*Sf9* insect cells (Thermo Fisher Scientific) were cultured in serum-free Ex-cell 420 medium (Sigma-Aldrich), and *Trichoplusnia ni* High Five™ (Thermo Fisher Scientific) were cultured in protein free ESF 921 insect cell culture media (Expression Systems LLC). Bacmid preparations, virus amplification in *Sf9*, and protein expression in *Trichoplusnia ni* High Five^*TM*^ insect cells was done according to standard protocols^63,64^.

Mature 20S CPs together with their assembly intermediates were purified by standard affinity purification on StrepTactin Sepharose High Performance (Cytiva) and subsequent SEC on a Superose6 Increase 10/300 (Cytiva) column in 25 mM HEPES pH 7.5 (KOH), 150 mM NaCl, 1 mM DTT.

### Structure determination by transmission electron cryo-microscopy

To prepare cryo-EM grids SEC peak fractions of mature 20S CPs were diluted to a concentration between 0.7 to 0.5 mg ml^-1^, SEC shoulder fractions containing 20S CP assembly intermediates were pooled, concentrated to 4 to 5 mg ml^-1^ and Fos-Cholin-8 was added to a final concentration of 0.25 mM. Quantifoil R1.2/1.3 Cu-200 grids (Quantifoil Micro Tools GmbH) were glow discharged for 30s in a PDC-32G-2 (Harrick Plasma Inc.), 3.5 μl samples were applied, and plunge frozen in liquid ethane/propane mix with a Vitribot Mark IV (Thermo Fisher Scientific) at 4°C and 100% humidity.

For cryo-EM data acquisition all grids were pre-screened for optimal particle distribution and ice thickness on an Glacios cryo-TEM (Thermo Fisher Scientific) operated at 200 kV equipped with a K2 Summit direct electron detector (DED) camera (Gatan). Data collection of the mature 20S CP dataset was carried out using a Glacios cryo-TEM. Data collection was set up with SerialEM version 4.1^65^ utilizing coma-corrected beam-image shift. One movie per hole was recorded in counting mode with a 3x3 multi hole record acquisition scheme at a pixel size of 1.181 Å/pixel with a nominal magnification of 36000x for the mature CP data set, or at pixel size of 1.885 Å/pixel with a nominal magnification of 22000x for the preholo/premature 20S CP data set. A total dose of 60 e-/Å^2^ was fractionated over 40 frames, with a target defocus range of -1.0 μm to -2.6 μm.

The data set for the 20S CP assembly intermediates was collected on a Titan Krios G2 cryo-TEM (Thermo Fisher Scientific) operated at 300 kV equipped with a Bio Quantum post-column energy filter (Gatan, 10eV) and K3 direct electron detector (DED) camera (Gatan). Data collection was set up with SerialEM version 4.1 utilizing coma-corrected beam-image shift. Three movies per hole were recorded with a 5x5 multi hole record acquisition scheme at a pixel size of 0.8512 Å/pixel with a nominal magnification of 105000x in counting mode (no CDS). A total dose of 68 e-/Å^2^ was fractionated over 30 frames, with a target defocus range of -1.0 μm to -2.6 μm.

Data processing was performed as follows. The Glacios K2 data set was processed with cryoSPARC version 4.2^66^, for details, see Fig. S1. The raw movies of the Titan Krios K3 dataset were on the fly motion corrected with FOCUS^67^, and motion corrected micrographs were imported into cryoSPARC for all subsequent processing steps (including patch CTF estimation, blob picking, template picking, 2D classification, heterogenous refinement, non-uniform refinement, manual sharpening, local resolution estimation, local filtering, and final post-processing with DeepEMhancer^68^), for details see Extended Data Fig. 2.

Model building and refinement was performed as follows. AlphaFold2^69,70^ models of PAC1-4 and POMP along with corresponding chains from a published model of the 20S CP (PDB 5LE5) were manually docked with ChimeraX version 1.5^71^ in all maps. Atomic models were locally adjusted in Coot version 0.9.8.7 ^72^ and iteratively refined with Phenix version 1.19.2^73^. We modelled into density when the trajectory of the backbone was unambiguous, and side-chains were placed when density was observed in manually sharpened or automatically sharpened map. There are additional regions of density around β2 in structure 1, β3 in structure 2, β4 in structure 3, β5 in structure 4 and β1/7 in structure 3. For which coordinates were not modelled due to ambiguity of these regions. For structure 1-5, the density for β2 CTE strand that grasps β3 partially overlaps with this element in the mature CP. However, due to a different trajectory for the preceding residues, we could not independently unambiguously assign the sequence as that in a mature 20S complex. Final refinement was done utilizing ISOLDE^74^ followed by Phenix refinement. For visualization figures were generated with UCSF ChimeraX, and Adobe Illustrator 2023.

## Acknowledgements

We thank Daniel Bollschweiler and Tillman Schäfer for maintaining the MPIB cryo-EM facility and assistance with data collection. We are grateful to J. Rajan Prabu for maintaining processing infrastructure and assistance with structural model building. This work was supported by Aligning Science Across Parkinsons (ASAP), the Max Planck Society for the Advancement of Science, and National Institutes of Health (NIH) grants R01-GM144367 (to J.H.), RO1-AG011085 (to J.W.H.). Michael J. Fox Foundation administers the grant ASAP-000282 on behalf of ASAP and itself. E.A.G. was the recipient of a postdoctoral fellowship from the Edward R. and Anne G. Lefler Center for the Study of Neurodegenerative Disorders.

## Author contributions

F.A. and B.A.S. designed research; F.A. contributed new reagents; F.A. and J.D. performed research; F.A., J.D., S.v.G., S.R., R.M.W., E.A.G. and J.W.H. analyzed data; J.W.H., J.H., and B.A.S. acquired funding; F.A., J.D., E.A.G., J.H., and B.A.S. wrote the manuscript with input from all authors.

### Data and Material Availability Statement

Cryo-EM maps have been deposited to the Electron Microscopy Data Bank (EMDB) with the following accession codes: map1: EMD-18755, map2: EMD-18757, map3: EMD-18758, map4: EMD-18773, map5: EMD-18759, preholo-20SCP: EMD-18761, premature-20SCP: EMD-18762, 20SCP: EMD-18760. Atomic coordinates have been deposited to RCSB Protein Data Bank (PDB) with the following accession codes: structure 1: PDB ID 8QYJ, structure 2: PDB ID 8QYL, structure 3: PDB ID 8QYM, structure 4: PDB ID 8QZ9, structure 5: PDB ID 8QYN, preholo-20SCP: PDB ID 8QYS, 20SCP: PDB ID 8QYO. Baculovirus transfer vectors for recombinant production of 20S core particles have been submitted to Addgene (# XXXX, #XXXX, #XXXX, #XXXX).

## Extended Data

**Ext. data figure 1.**
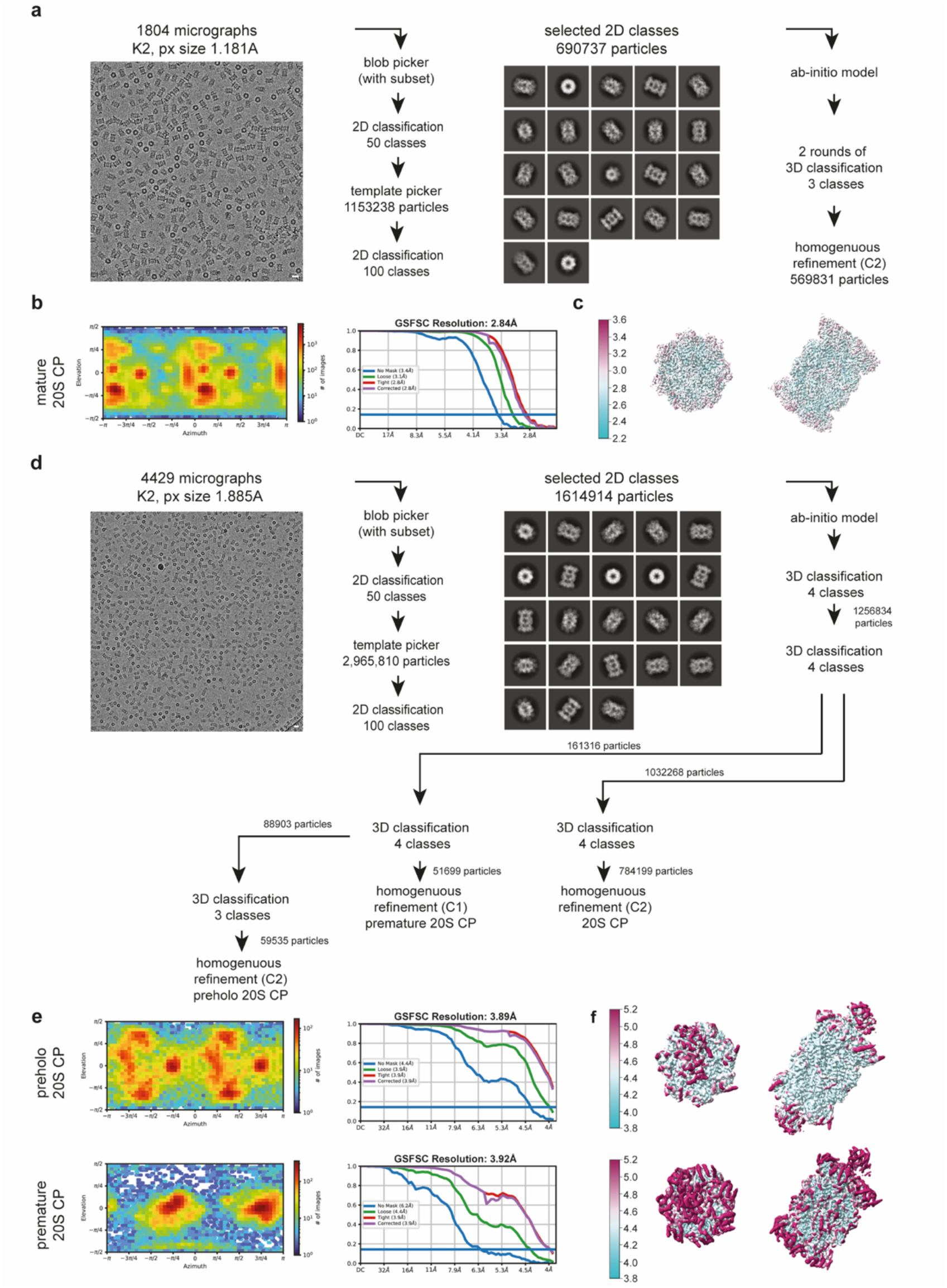
Structure determination of recombinant human 20S proteasome. **a**, Cryo-EM image processing scheme for mature human 20S proteasome. **b**, Angular distribution and GSFSC curves of human 20S proteasome map. **c**, Local resolution of the cryo-EM map of human 20S proteasome map. **d**, Cryo-EM image processing scheme for preholo 20S CP and premature 20S CP. **e**, Angular distribution and GSFSC curves of preholo 20S CP (top) and premature 20S CP (bottom) maps. **f**, Local resolution of the cryo-EM maps of preholo 20S CP (top) and premature 20S CP (bottom) Scale bar in all micrographs represents 15 nm.

**Ext. data figure 2.**
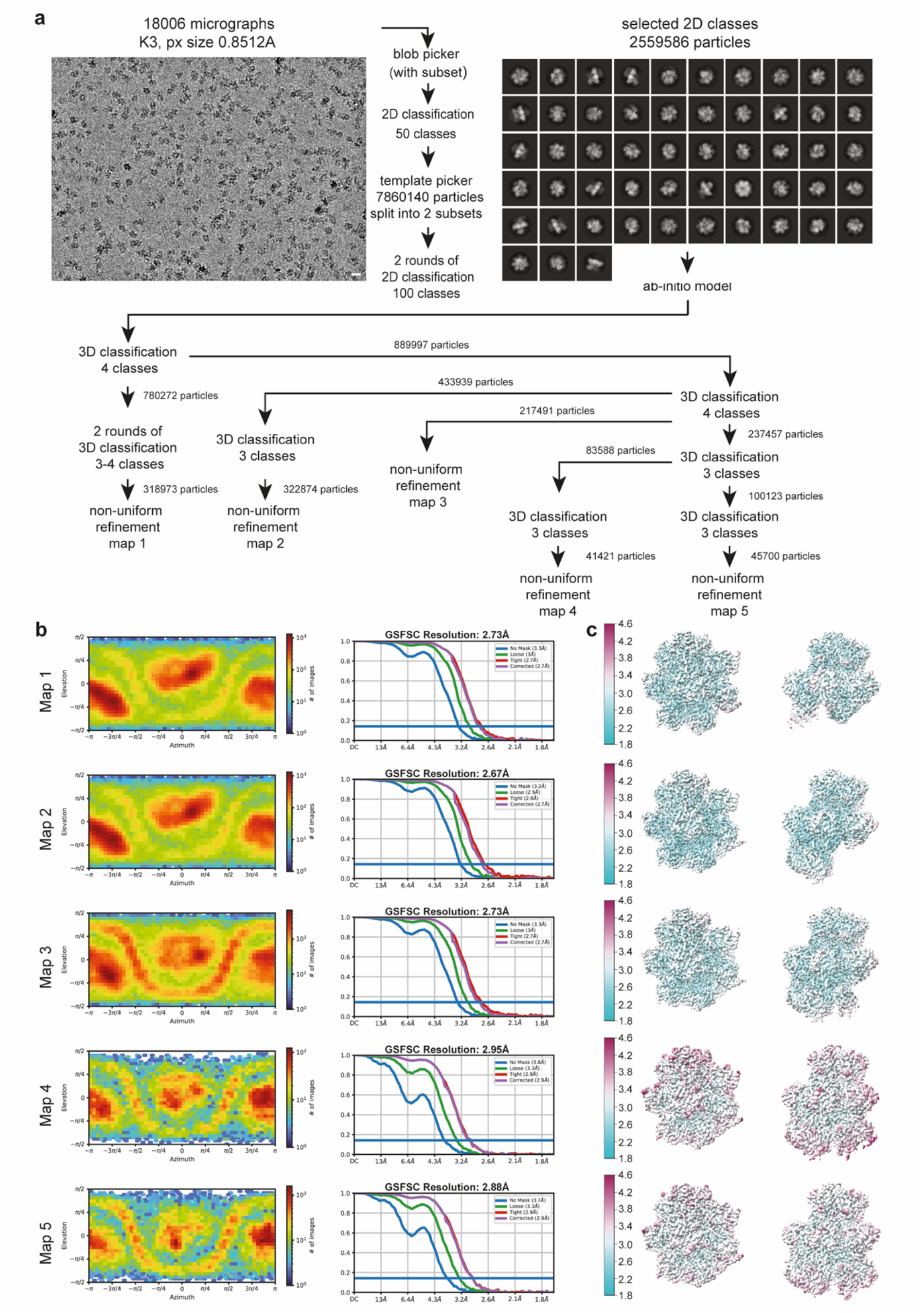
Structure determination of recombinant human proteasome assembly intermediates. **a**, Cryo-EM image processing scheme for the human proteasome assembly intermediates. **b**, Angular distribution and GSFSC curves of human proteasome assembly intermediates. **c**, Local resolution of the cryo-EM maps of human proteasome assembly intermediates. Scale bar in all micrographs represents 15 nm.

**Ext. data figure 3.**
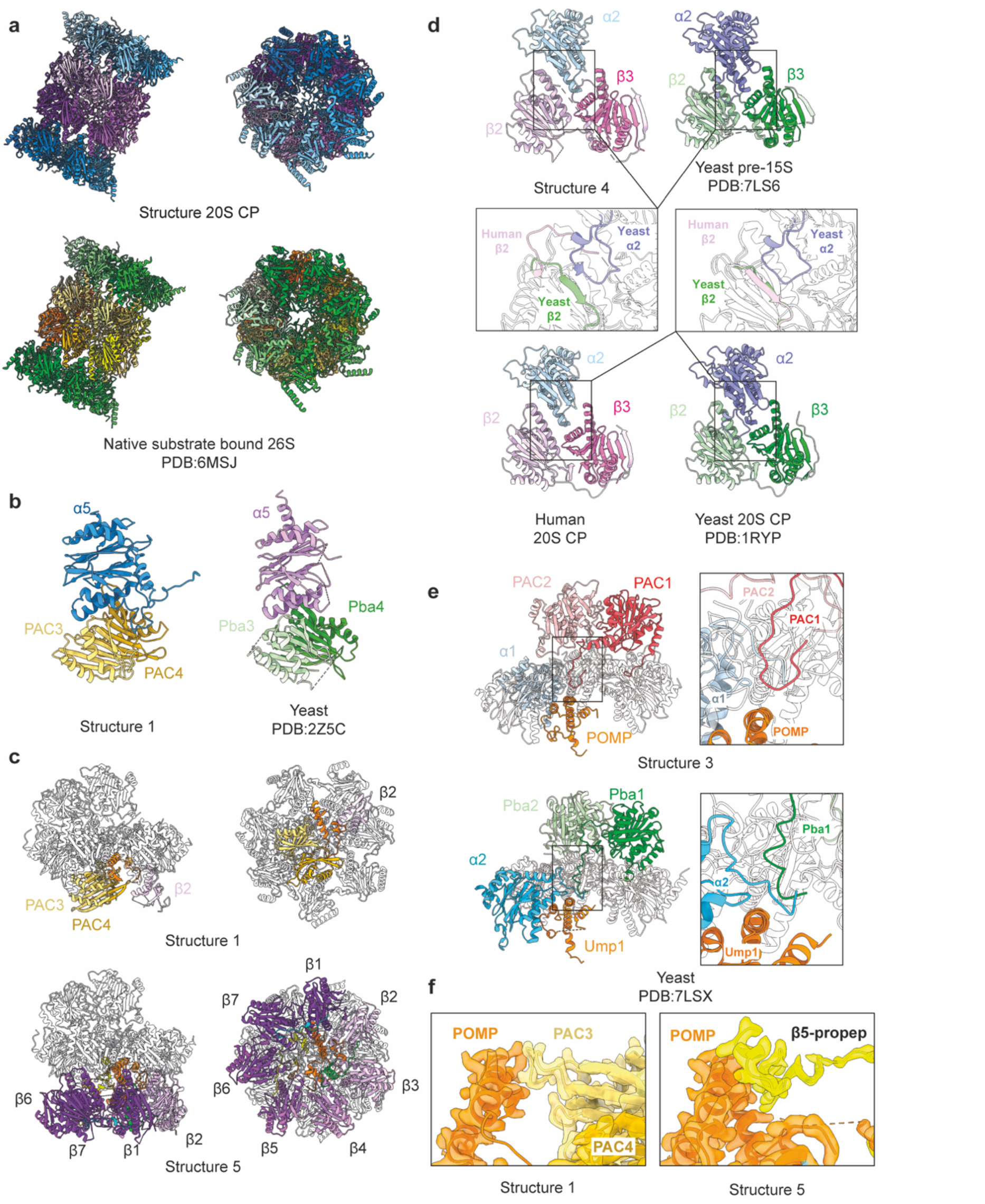
Structural comparison of human 20SCP assembly intermediates with yeast orthologs. **a**, Comparison of human 20S core particle with native substrate bound 26S proteasome (PDB: 6MSJ, only 20S core particle is shown). **b**, Comparison of human PAC3/4 (yellow) with yeast Pba3/4 (green, PDB:2Z5C) reveals their structural similarity. **c**, Comparison of structure 1 and structure 5 shows the position of PAC3/4 in structure 1 is occupied by β4-7 later in structure 5. **d**, Comparison of human β2 (pink) with yeast β2 (green, PDB:7LS6/1RYP) in pre-15S (top) and mature 20S CP (bottom) shows the insertion of yeast α2 changes the trajectory of yeast β2 C-terminal loop compared to human in immature 20S CP. **e**, Comparison of human PAC1/2 (red) with yeast Pba1/2 (green, PDB:7LSX) shows the N-terminus of PAC1 and Pba1 interacts with human α1 and yeast α2, respectively, and make close contact with POMP/Ump1. **f**, Comparison of structure 1 POMP interaction with PAC3 and structure 5 POMP interaction with β5 propeptide

**Ext. data figure 4.**
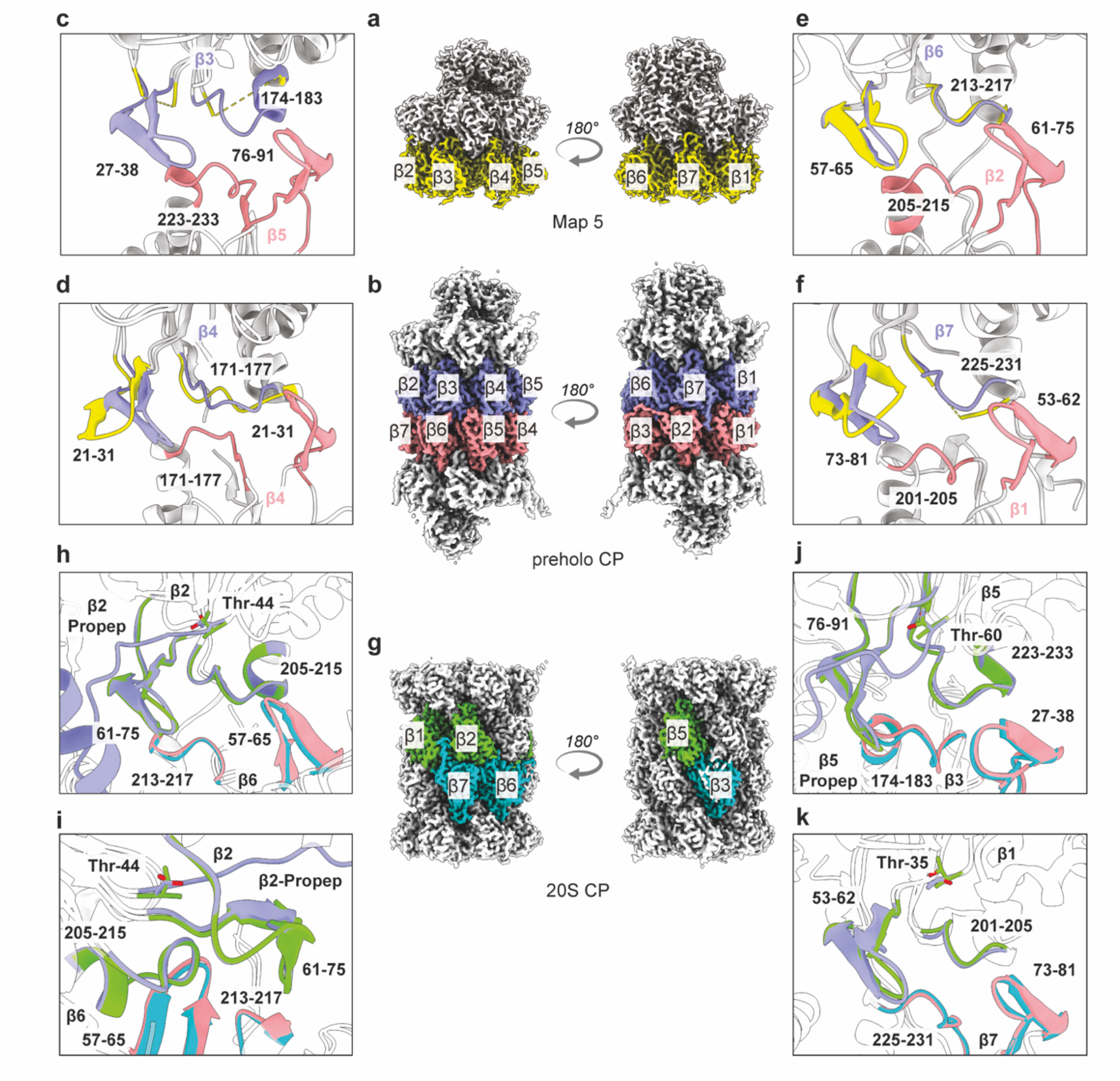
Fusion-tetrad formed by the opposing β subunits upon half CP fusion. **a**, Cryo-EM map of structure 5 with β-subunits colored in yellow. **b**, Cryo-EM map of preholo CP with β-subunits of one half CP colored in purple and the other half CP colored in salmon. **c-f**, Close-up view of fusion-tetrad between β3 (structure 5 in yellow and preholo 20SCP in purple) and β5 from the opposite half CP (salmon) (c), two opposing β4 (d), β6 and β2 from the opposite half CP (e) and β7 and β1 from the opposite half CP (f). **g**, Cryo-EM map of mature 20S CP with β-subunits of one half CP colored in green and the other half CP colored in blue. **h-k**, Close-up view of fusion-tetrad between β2 (preholo 20SCP in purple and mature 20SCP in green) and β5 from the opposite half CP (preholo 20SCP in salmon and mature 20SCP in blue) (h-i), β5 and β3 from the opposite half CP (j), and β1 and β7 from the opposite half CP (k).

**Ext. data figure 5.**
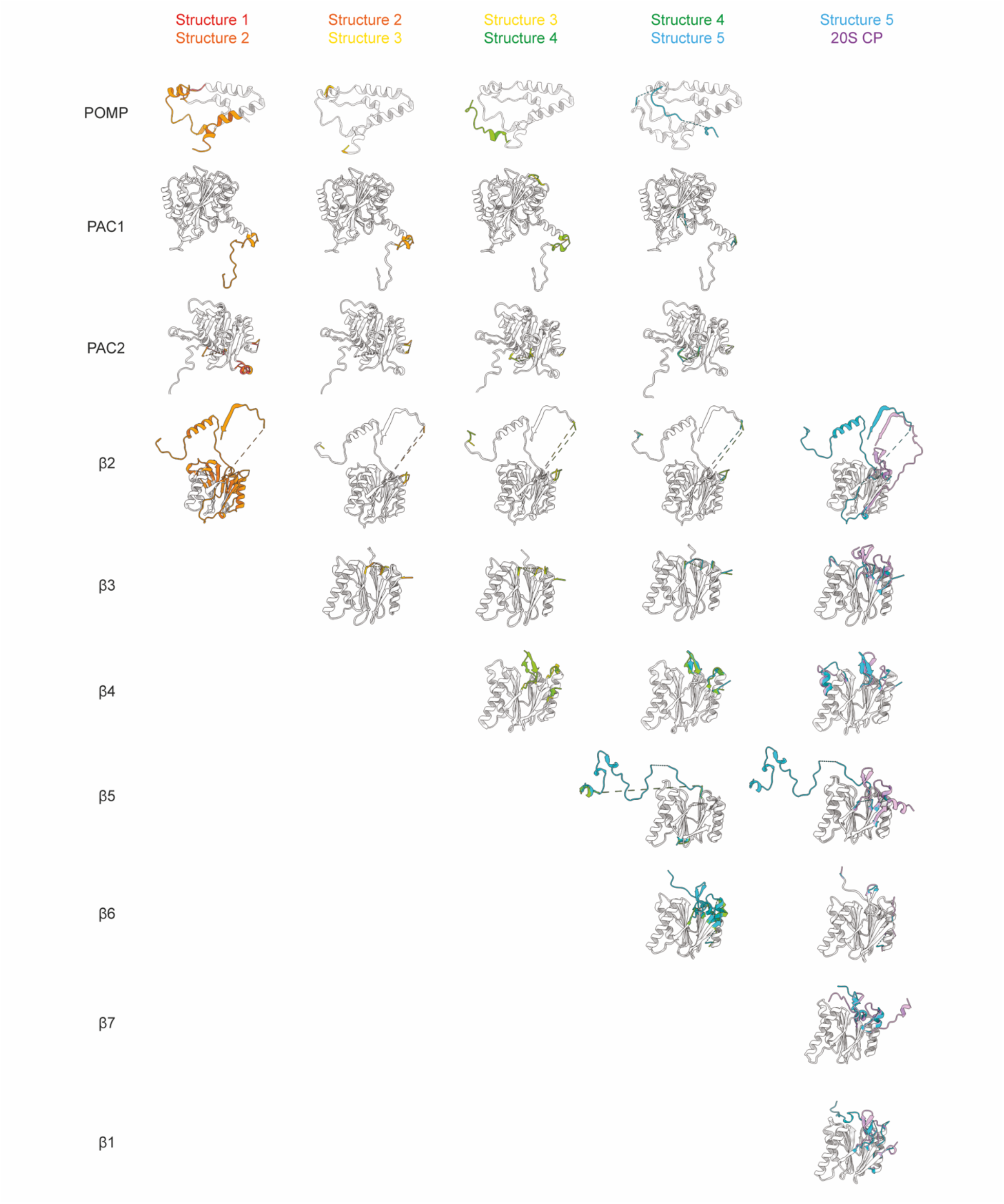
Structures of assembly chaperons and β subunits across β-ring formation.

**Ext. data figure 6.**
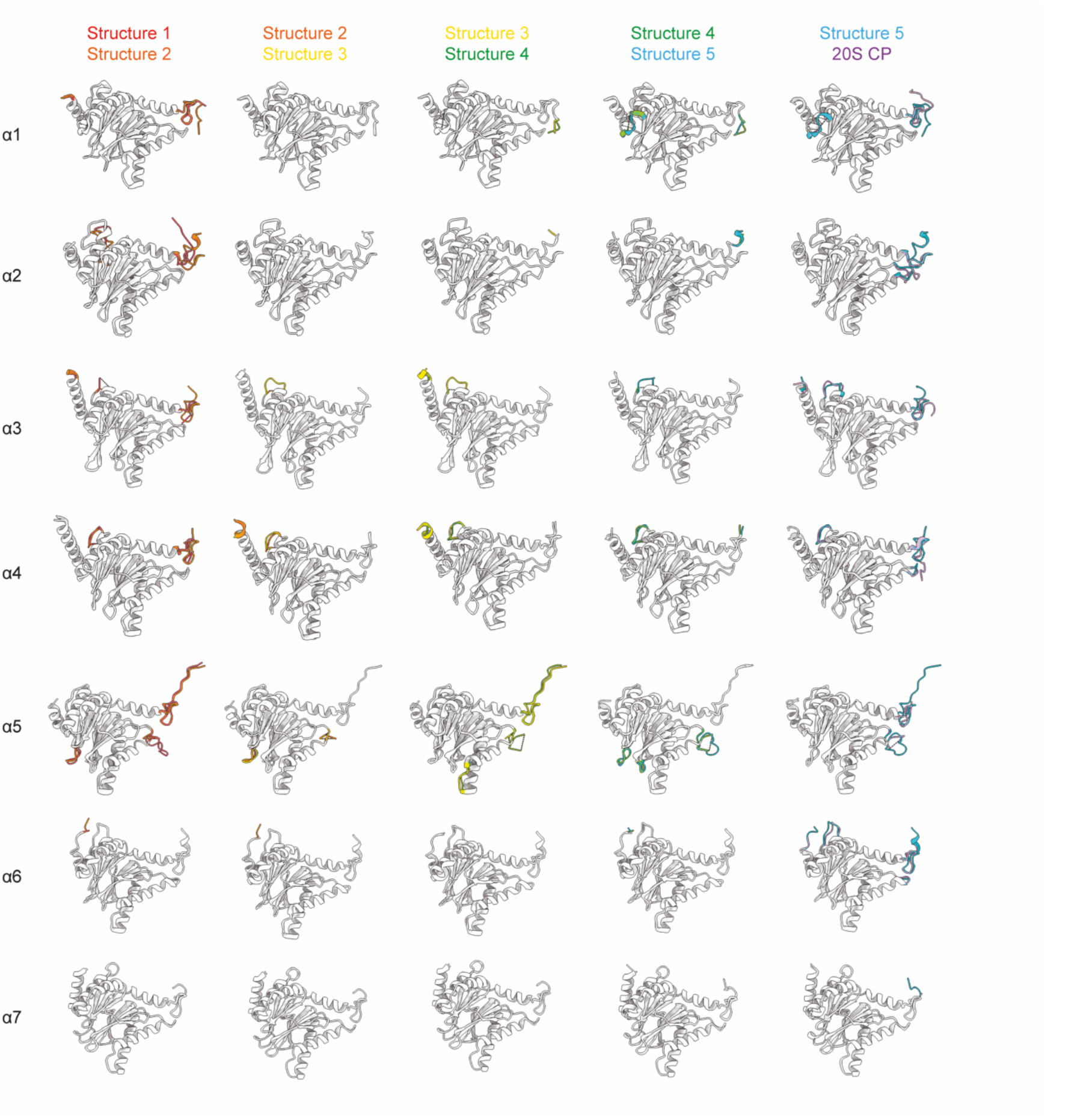
Structural comparison of α subunits across β-ring formation.

## References

1. Lowe, J., Stock, D., Jap, B., Zwickl, P., Baumeister, W. & Huber, R. Crystal structure of the 20S proteasome from the archaeon T. acidophilum at 3.4 A resolution. Science 268, 533–539 (1995). 10.1126/science.7725097

2. Groll, M. et al. Structure of 20S proteasome from yeast at 2.4 A resolution. Nature 386, 463–471 (1997). 10.1038/386463a0

3. Groll, M. et al. A gated channel into the proteasome core particle. Nat Struct Biol 7, 1062–1067 (2000). 10.1038/80992

4. Unno, M. et al. The structure of the mammalian 20S proteasome at 2.75 A resolution. Structure 10, 609–618 (2002). 10.1016/s0969-2126(02)00748-7

5. Arendt, C. S. & Hochstrasser, M. Identification of the yeast 20S proteasome catalytic centers and subunit interactions required for active-site formation. Proc Natl Acad Sci U S A 94, 7156–7161 (1997). 10.1073/pnas.94.14.7156

6. Chen, X., Htet, Z. M., Lopez-Alfonzo, E., Martin, A. & Walters, K. J. Proteasome interaction with ubiquitinated substrates: from mechanisms to therapies. FEBS J 288, 5231–5251 (2021). 10.1111/febs.15638

7. Whitby, F. G. et al. Structural basis for the activation of 20S proteasomes by 11S regulators. Nature 408, 115–120 (2000). 10.1038/35040607

8. Smith, D. M., Chang, S. C., Park, S., Finley, D., Cheng, Y. & Goldberg, A. L. Docking of the proteasomal ATPases’ carboxyl termini in the 20S proteasome’s alpha ring opens the gate for substrate entry. Mol Cell 27, 731–744 (2007). 10.1016/j.molcel.2007.06.033

9. Smith, D. M., Kafri, G., Cheng, Y., Ng, D., Walz, T. & Goldberg, A. L. ATP binding to PAN or the 26S ATPases causes association with the 20S proteasome, gate opening, and translocation of unfolded proteins. Mol Cell 20, 687–698 (2005). 10.1016/j.molcel.2005.10.019

10. Rabl, J., Smith, D. M., Yu, Y., Chang, S. C., Goldberg, A. L. & Cheng, Y. Mechanism of gate opening in the 20S proteasome by the proteasomal ATPases. Mol Cell 30, 360–368 (2008). 10.1016/j.molcel.2008.03.004

11. Kohler, A., Cascio, P., Leggett, D. S., Woo, K. M., Goldberg, A. L. & Finley, D. The axial channel of the proteasome core particle is gated by the Rpt2 ATPase and controls both substrate entry and product release. Mol Cell 7, 1143–1152 (2001). 10.1016/s1097-2765(01)00274-x

12. Wehmer, M. et al. Structural insights into the functional cycle of the ATPase module of the 26S proteasome. Proc Natl Acad Sci U S A 114, 1305–1310 (2017). 10.1073/pnas.1621129114

13. de la Pena, A. H., Goodall, E. A., Gates, S. N., Lander, G. C. & Martin, A. Substrate-engaged 26S proteasome structures reveal mechanisms for ATP-hydrolysis-driven translocation. Science 362 (2018). 10.1126/science.aav0725

14. Forster, A., Whitby, F. G. & Hill, C. P. The pore of activated 20S proteasomes has an ordered 7-fold symmetric conformation. EMBO J 22, 4356–4364 (2003). 10.1093/emboj/cdg436

15. Sadre-Bazzaz, K., Whitby, F. G., Robinson, H., Formosa, T. & Hill, C. P. Structure of a Blm10 complex reveals common mechanisms for proteasome binding and gate opening. Mol Cell 37, 728–735 (2010). 10.1016/j.molcel.2010.02.002

16. Toste Rego, A. & da Fonseca, P. C. A. Characterization of Fully Recombinant Human 20S and 20S-PA200 Proteasome Complexes. Mol Cell 76, 138–147 e135 (2019). 10.1016/j.molcel.2019.07.014

17. Guan, H. et al. Cryo-EM structures of the human PA200 and PA200-20S complex reveal regulation of proteasome gate opening and two PA200 apertures. PLoS Biol 18, e3000654 (2020). 10.1371/journal.pbio.3000654

18. Itzhak, D. N., Tyanova, S., Cox, J. & Borner, G. H. Global, quantitative and dynamic mapping of protein subcellular localization. Elife 5 (2016). 10.7554/eLife.16950

19. Lehrbach, N. J., Breen, P. C. & Ruvkun, G. Protein Sequence Editing of SKN-1A/Nrf1 by Peptide:N-Glycanase Controls Proteasome Gene Expression. Cell 177, 737–750 e715 (2019). 10.1016/j.cell.2019.03.035

20. Watanabe, A., Yashiroda, H., Ishihara, S., Lo, M. & Murata, S. The Molecular Mechanisms Governing the Assembly of the Immuno- and Thymoproteasomes in the Presence of Constitutive Proteasomes. Cells 11 (2022). 10.3390/cells11091580

21. Schnell, H. M., Walsh, R. M., Rawson, S. & Hanna, J. Chaperone-mediated assembly of the proteasome core particle - recent developments and structural insights. J Cell Sci 135 (2022). 10.1242/jcs.259622

22. Rousseau, A. & Bertolotti, A. Regulation of proteasome assembly and activity in health and disease. Nat Rev Mol Cell Biol 19, 697–712 (2018). 10.1038/s41580-018-0040-z

23. Burri, L., Hockendorff, J., Boehm, U., Klamp, T., Dohmen, R. J. & Levy, F. Identification and characterization of a mammalian protein interacting with 20S proteasome precursors. Proc Natl Acad Sci U S A 97, 10348–10353 (2000). 10.1073/pnas.190268597

24. Griffin, T. A., Slack, J. P., McCluskey, T. S., Monaco, J. J. & Colbert, R. A. Identification of proteassemblin, a mammalian homologue of the yeast protein, Ump1p, that is required for normal proteasome assembly. Mol Cell Biol Res Commun 3, 212–217 (2000). 10.1006/mcbr.2000.0213

25. Sa-Moura, B. et al. Biochemical and biophysical characterization of recombinant yeast proteasome maturation factor ump1. Comput Struct Biotechnol J 7, e201304006 (2013). 10.5936/csbj.201304006

26. Witt, E., Zantopf, D., Schmidt, M., Kraft, R., Kloetzel, P. M. & Kruger, E. Characterisation of the newly identified human Ump1 homologue POMP and analysis of LMP7(beta 5i) incorporation into 20 S proteasomes. J Mol Biol 301, 1–9 (2000). 10.1006/jmbi.2000.3959

27. Hirano, Y. et al. A heterodimeric complex that promotes the assembly of mammalian 20S proteasomes. Nature 437, 1381–1385 (2005). 10.1038/nature04106

28. Hirano, Y. et al. Cooperation of multiple chaperones required for the assembly of mammalian 20S proteasomes. Mol Cell 24, 977–984 (2006). 10.1016/j.molcel.2006.11.015

29. Le Tallec, B., Barrault, M. B., Courbeyrette, R., Guerois, R., Marsolier-Kergoat, M. C. & Peyroche, A. 20S proteasome assembly is orchestrated by two distinct pairs of chaperones in yeast and in mammals. Mol Cell 27, 660–674 (2007). 10.1016/j.molcel.2007.06.025

30. Kusmierczyk, A. R., Kunjappu, M. J., Funakoshi, M. & Hochstrasser, M. A multimeric assembly factor controls the formation of alternative 20S proteasomes. Nat Struct Mol Biol 15, 237–244 (2008). 10.1038/nsmb.1389

31. Chen, P. & Hochstrasser, M. Autocatalytic subunit processing couples active site formation in the 20S proteasome to completion of assembly. Cell 86, 961–972 (1996). 10.1016/s0092-8674(00)80171-3

32. Jager, S., Groll, M., Huber, R., Wolf, D. H. & Heinemeyer, W. Proteasome beta-type subunits: unequal roles of propeptides in core particle maturation and a hierarchy of active site function. J Mol Biol 291, 997–1013 (1999). 10.1006/jmbi.1999.2995

33. Hirano, Y. et al. Dissecting beta-ring assembly pathway of the mammalian 20S proteasome. EMBO J 27, 2204–2213 (2008). 10.1038/emboj.2008.148

34. Frentzel, S., Pesold-Hurt, B., Seelig, A. & Kloetzel, P. M. 20 S proteasomes are assembled via distinct precursor complexes. Processing of LMP2 and LMP7 proproteins takes place in 13-16 S preproteasome complexes. J Mol Biol 236, 975–981 (1994). 10.1016/0022-2836(94)90003-5

35. Dahlqvist, J. et al. A single-nucleotide deletion in the POMP 5’ UTR causes a transcriptional switch and altered epidermal proteasome distribution in KLICK genodermatosis. Am J Hum Genet 86, 596–603 (2010). 10.1016/j.ajhg.2010.02.018

36. Poli, M. C. et al. Heterozygous Truncating Variants in POMP Escape Nonsense-Mediated Decay and Cause a Unique Immune Dysregulatory Syndrome. Am J Hum Genet 102, 1126–1142 (2018). 10.1016/j.ajhg.2018.04.010

37. de Jesus, A. A. et al. Novel proteasome assembly chaperone mutations in PSMG2/PAC2 cause the autoinflammatory interferonopathy CANDLE/PRAAS4. J Allergy Clin Immunol 143, 1939–1943 e1938 (2019). 10.1016/j.jaci.2018.12.1012

38. Sasaki, K. et al. PAC1 gene knockout reveals an essential role of chaperone-mediated 20S proteasome biogenesis and latent 20S proteasomes in cellular homeostasis. Mol Cell Biol 30, 3864–3874 (2010). 10.1128/MCB.00216-10

39. Yashiroda, H. et al. Crystal structure of a chaperone complex that contributes to the assembly of yeast 20S proteasomes. Nat Struct Mol Biol 15, 228–236 (2008). 10.1038/nsmb.1386

40. Wu, W. et al. PAC1-PAC2 proteasome assembly chaperone retains the core alpha4-alpha7 assembly intermediates in the cytoplasm. Genes Cells 23, 839–848 (2018). 10.1111/gtc.12631

41. Stadtmueller, B. M. et al. Structure of a proteasome Pba1-Pba2 complex: implications for proteasome assembly, activation, and biological function. J Biol Chem 287, 37371–37382 (2012). 10.1074/jbc.M112.367003

42. Takagi, K. et al. Pba3-Pba4 heterodimer acts as a molecular matchmaker in proteasome alpha-ring formation. Biochem Biophys Res Commun 450, 1110–1114 (2014). 10.1016/j.bbrc.2014.06.119

43. Seemuller, E., Lupas, A. & Baumeister, W. Autocatalytic processing of the 20S proteasome. Nature 382, 468–471 (1996). 10.1038/382468a0

44. Huber, E. M., Heinemeyer, W., Li, X., Arendt, C. S., Hochstrasser, M. & Groll, M. A unified mechanism for proteolysis and autocatalytic activation in the 20S proteasome. Nat Commun 7, 10900 (2016). 10.1038/ncomms10900

45. Schmidtke, G., Schmidt, M. & Kloetzel, P. M. Maturation of mammalian 20 S proteasome: purification and characterization of 13 S and 16 S proteasome precursor complexes. J Mol Biol 268, 95–106 (1997). 10.1006/jmbi.1997.0947

46. Nandi, D., Woodward, E., Ginsburg, D. B. & Monaco, J. J. Intermediates in the formation of mouse 20S proteasomes: implications for the assembly of precursor beta subunits. EMBO J 16, 5363–5375 (1997). 10.1093/emboj/16.17.5363

47. Li, X., Kusmierczyk, A. R., Wong, P., Emili, A. & Hochstrasser, M. beta-Subunit appendages promote 20S proteasome assembly by overcoming an Ump1-dependent checkpoint. EMBO J 26, 2339–2349 (2007). 10.1038/sj.emboj.7601681

48. Schnell, H. M. et al. Structures of chaperone-associated assembly intermediates reveal coordinated mechanisms of proteasome biogenesis. Nat Struct Mol Biol 28, 418–425 (2021). 10.1038/s41594-021-00583-9

49. Groll, M. et al. The catalytic sites of 20S proteasomes and their role in subunit maturation: a mutational and crystallographic study. Proc Natl Acad Sci U S A 96, 10976–10983 (1999). 10.1073/pnas.96.20.10976

50. Beckwith, R., Estrin, E., Worden, E. J. & Martin, A. Reconstitution of the 26S proteasome reveals functional asymmetries in its AAA+ unfoldase. Nat Struct Mol Biol 20, 1164–1172 (2013). 10.1038/nsmb.2659

51. Lander, G. C., Estrin, E., Matyskiela, M. E., Bashore, C., Nogales, E. & Martin, A. Complete subunit architecture of the proteasome regulatory particle. Nature 482, 186–191 (2012). 10.1038/nature10774

52. Silhan, J. et al. Structural elucidation of recombinant Trichomonas vaginalis 20S proteasome bound to covalent inhibitors. bioRxiv (2023). 10.1101/2023.08.17.553660

53. Weissmann, F. et al. biGBac enables rapid gene assembly for the expression of large multisubunit protein complexes. Proc Natl Acad Sci U S A 113, E2564–2569 (2016). 10.1073/pnas.1604935113

54. Vijayachandran, L. S. et al. Robots, pipelines, polyproteins: enabling multiprotein expression in prokaryotic and eukaryotic cells. J Struct Biol 175, 198–208 (2011). 10.1016/j.jsb.2011.03.007

55. Dong, Y. et al. Cryo-EM structures and dynamics of substrate-engaged human 26S proteasome. Nature 565, 49–55 (2019). 10.1038/s41586-018-0736-4

56. Schnell, H. M., Ang, J., Rawson, S., Walsh, R. M., Jr., Micoogullari, Y. & Hanna, J. Mechanism of proteasome gate modulation by assembly chaperones Pba1 and Pba2. J Biol Chem 298, 101906 (2022). 10.1016/j.jbc.2022.101906

57. Kurimoto, E. et al. Crystal structure of human proteasome assembly chaperone PAC4 involved in proteasome formation. Protein Sci 26, 1080–1085 (2017). 10.1002/pro.3153

58. Satoh, T., Yagi-Utsumi, M., Okamoto, K., Kurimoto, E., Tanaka, K. & Kato, K. Molecular and Structural Basis of the Proteasome alpha Subunit Assembly Mechanism Mediated by the Proteasome-Assembling Chaperone PAC3-PAC4 Heterodimer. Int J Mol Sci 20 (2019). 10.3390/ijms20092231

59. Fricke, B., Heink, S., Steffen, J., Kloetzel, P. M. & Kruger, E. The proteasome maturation protein POMP facilitates major steps of 20S proteasome formation at the endoplasmic reticulum. EMBO Rep 8, 1170–1175 (2007). 10.1038/sj.embor.7401091

60. Ramos, P. C., Marques, A. J., London, M. K. & Dohmen, R. J. Role of C-terminal extensions of subunits beta2 and beta7 in assembly and activity of eukaryotic proteasomes. J Biol Chem 279, 14323–14330 (2004). 10.1074/jbc.M308757200

61. Walsh, R. M., Jr., Rawson, S., Schnell, H. M., Velez, B., Rajakumar, T. & Hanna, J. Structure of the preholoproteasome reveals late steps in proteasome core particle biogenesis. Nat Struct Mol Biol (2023). 10.1038/s41594-023-01081-w

62. Leggett, D. S. et al. Multiple associated proteins regulate proteasome structure and function. Mol Cell 10, 495–507 (2002). 10.1016/s1097-2765(02)00638-x

63. Fitzgerald, D. J., Berger, P., Schaffitzel, C., Yamada, K., Richmond, T. J. & Berger, I. Protein complex expression by using multigene baculoviral vectors. Nat Methods 3, 1021–1032 (2006). 10.1038/nmeth983

64. Bieniossek, C., Richmond, T. J. & Berger, I. MultiBac: multigene baculovirus-based eukaryotic protein complex production. Curr Protoc Protein Sci Chapter 5, Unit 5 20 (2008). 10.1002/0471140864.ps0520s51

65. Mastronarde, D. N. Automated electron microscope tomography using robust prediction of specimen movements. J Struct Biol 152, 36–51 (2005). 10.1016/j.jsb.2005.07.007

66. Punjani, A., Rubinstein, J. L., Fleet, D. J. & Brubaker, M. A. cryoSPARC: algorithms for rapid unsupervised cryo-EM structure determination. Nat Methods 14, 290–296 (2017). 10.1038/nmeth.4169

67. Biyani, N. et al. Focus: The interface between data collection and data processing in cryo-EM. J Struct Biol 198, 124–133 (2017). 10.1016/j.jsb.2017.03.007

68. Sanchez-Garcia, R., Gomez-Blanco, J., Cuervo, A., Carazo, J. M., Sorzano, C. O. S. & Vargas, J. DeepEMhancer: a deep learning solution for cryo-EM volume post-processing. Commun Biol 4, 874 (2021). 10.1038/s42003-021-02399-1

69. Jumper, J. et al. Highly accurate protein structure prediction with AlphaFold. Nature 596, 583–589 (2021). 10.1038/s41586-021-03819-2

70. Jumper, J. & Hassabis, D. Protein structure predictions to atomic accuracy with AlphaFold. Nat Methods 19, 11–12 (2022). 10.1038/s41592-021-01362-6

71. Goddard, T. D. et al. UCSF ChimeraX: Meeting modern challenges in visualization and analysis. Protein Sci 27, 14–25 (2018). 10.1002/pro.3235

72. Emsley, P., Lohkamp, B., Scott, W. G. & Cowtan, K. Features and development of Coot. Acta Crystallogr D Biol Crystallogr 66, 486–501 (2010). 10.1107/S0907440007493

73. Liebschner, D. et al. Macromolecular structure determination using X-rays, neutrons and electrons: recent developments in Phenix. Acta Crystallogr D Struct Biol 75, 861–877 (2019). 10.1107/S2059798319011471

74. Croll, T. I. ISOLDE: a physically realistic environment for model building into low-resolution electron-density maps. Acta Crystallogr D Struct Biol 74, 519–530 (2018). 10.1107/S2059798318002425

